# Evidence of a novel viral membrane fusion mechanism shared by the Hepaci, Pegi and Pestiviruses

**DOI:** 10.1101/2022.10.18.512720

**Authors:** Michael R. Oliver, Kamilla Toon, Charlotte B. Lewis, Stephen Devlin, Robert J. Gifford, Joe Grove

## Abstract

Enveloped viruses encode specialised glycoproteins that mediate fusion of viral and host membranes. Discovery and understanding of the molecular mechanisms of fusion has been achieved through structural analyses of glycoproteins from many different viruses, and yet the fusion mechanisms of some viral genera remain unknown. We have employed systematic genome annotation and AlphaFold modelling to predict the structures of the E1E2 glycoproteins from sixty viral species in the Hepaci, Pegi and Pestivirus genera. Whilst the predicted structure of E2 varied widely, E1 exhibited a very consistent fold across genera, despite little or no homology at the sequence level. Critically, the structure of E1 is unlike any other known viral glycoprotein. This is the first evidence that the Hepaci, Pegi and Pestiviruses possess a common and novel membrane fusion mechanism. Comparison of E1E2 models from various species reveals recurrent features that are likely to be mechanistically important and sheds light on the evolution of membrane fusion in these viral genera. These findings provide new fundamental understanding of viral membrane fusion and are relevant to structure-guided vaccinology.

## Background

Enveloped viruses encode specialised glycoproteins that undergo dramatic conformational change to mediate fusion of viral and host membranes. To date, three distinct classes of fusion protein are known. Class-I, as found in, for example, influenzaviruses, retroviruses, coronaviruses ^1–7^; class-II, in flaviviruses, alphaviruses ^8–12^; class-III, in rhabdoviruses, herpesviruses, baculoviruses ^13–16^. Mechanistic conservation suggests each class of fusion protein arose from ancient progenitors that underwent genetic transfer within the virome. However, subsequent evolutionary divergence has eliminated any genetic homology between the glycoproteins within a given class. Therefore, mechanistic/evolutionary grouping of fusion proteins cannot be inferred from primary sequence and is only revealed by structural analysis. Moreover, determination of the structure of viral glycoproteins in both their pre- and post-fusion states is critical to understanding the conformational rearrangements that drive membrane fusion. This knowledge has informed vaccinology ^17–19^ and provided insights on fusion in eukaryotes ^20^

There remain genera of viruses for which the mechanism of fusion remains unknown; for example, the Hepaciviruses (e.g. hepatitis C virus, HCV-C), Pegiviruses and Pestiviruses (e.g. bovine viral diarrhoea virus, BVDV), which are all members of the Flaviviridae family. Whilst classical Flaviviruses (e.g. dengue virus) possess class-II fusion proteins, the current structural understanding of HCV and BVDV suggest distinct and, hitherto, unknown fusion mechanism(s) ^21–24^. Hepaci, Pegi and Pestiviruses achieve fusion via minor (E1) and major (E2) glycoproteins that act in concert. HCV and BVDV E2 are structurally distinct, raising the possibility that they have different fusion mechanisms. Here, we propose a common fusion mechanism across the Hepaci, Pegi and Pestiviruses. This is evidenced by high structural conservation in the minor glycoprotein E1, revealed by *ab initio* protein structure prediction. Our comparison of E1E2 models from varied species provide insights on mechanism and evolutionary origin.

## Structural conservation of Hepaci, Pegi and Pestivirus E1

We performed systematic genome alignment and annotation to generate matched E1E2 targets for AlphaFold protein structure prediction using the ColabFold platform ^25,26^ (total of 61 E1E2 targets), yielding high confidence E1E2 dimer models from 11, 3 and 10 Hepaci, Pegi and Pestiviruses respectively (see Supplementary Materials for benchmarking performance, Fig. S1, and a detailed description of AlphaFold modelling, Fig. S2-S14). E2 models were structurally distinct between (and, to some extent, within) viral genera, whereas E1 models exhibited high structural similarity (Fig. 1A). These observations were supported by unbiased all-against-all analysis of structural similarity using the DALI server ^27^ (Fig. 1B & Fig. S15). E1 exhibits three conserved structural motifs found across the viral genera Fig. 1C: i) a central anti-parallel beta-sheet ii) a helical hairpin iii) a transmembrane proximal region.

**Figure 1.**
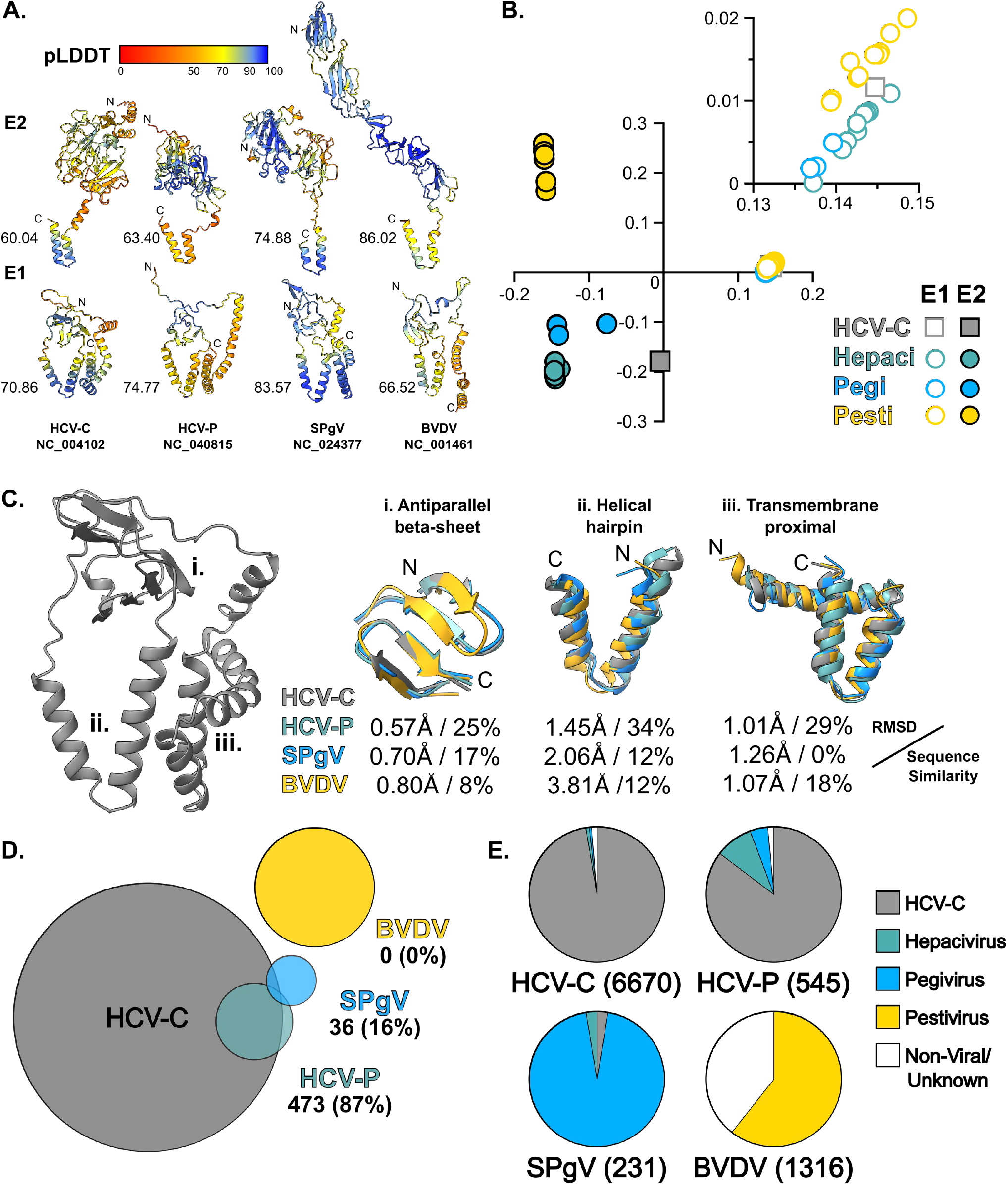
Structural conservation of E1 across the Hepaci, Pegi and Pestiviruses. **A.** AlphaFold predicted E1 and E2 structures, shown as ribbon diagrams, for Hepatitis C Virus (HCV-C), Hepacivirus-P (HCV-P), Simian Pegivirus (SPgV) and Bovine Viral Diarrhoea Virus (BVDV). Models are colour coded by pLDDT prediction confidence. **B.** Comparison and clustering of E1 and E2 (from 24 species) based on structural similarity through correspondence analysis, the plot is a projection of the first two eigenvectors. Inset axes are enlarged to highlight the E1 structural cluster. **C.** Consistent features of E1 shared across genera. Left, structure of HCV-C E1 for context. Ribbon diagrams of overlaid features for HCV-C, HCV-P, SPgV and BVDV, are accompanied by root mean square deviation (RMSD) and sequence similarity values by comparison to HCV-C. **D.** Venn diagram demonstrating overlap in underlying multiple sequence alignment (MSA) data used to generate predicted structures for HCV-C, HCV-P, SPgV and BVDV. The number of sequences (and percentage) overlapping with the HCV-C MSA is provided. **E.** The constituent sequences from each MSA, classified by source.The number of sequences in each MSA is provided in parenthesis.

AlphaFold/ColabFold automatically generates multiple sequence alignments based on homology to the given target sequence ^25,26^; protein structure is inferred using the evolutionary relationships revealed in these alignments. Therefore, structural similarity between targets can be simply driven by primary sequence homology (i.e. two closely related target proteins are likely to yield similar structures). Consequently, agreement in structure prediction between multiple homologous targets is not a guarantee of accuracy; they could all be similarly incorrect. However, if targets with little/no homology yield similar structures there is more confidence in their veracity. We, therefore, compared the sequence data underlying our various E1E2 structure predictions. Hepacivirus E1E2 models draw heavily on HCV-C sequences, however, with greater genetic distance the structural models become increasingly independent. Pestivirus E1E2 models share no overlap with Hepaci or Pegiviruses in their underlying sequence data (Fig. 1D & E). Therefore, Hepaci/Pegi and Pestivirus E1E2 models are completely independent, providing high confidence that their apparent structural similarity is accurate. The high degree of E1 structural homology across these genetically divergent viral genera is the first direct evidence that they share a mechanistically conserved fusion mechanism.

## Molecular architecture of E1

The predicted structure of E1 exhibits a conserved topology and compact architecture in close apposition to the viral membrane, the approximate location of which is inferred by the position of the E1 transmembrane domain (Fig. 2A & B). Towards the N-terminus a central beta-sheet sits on top of a distinctive helical hairpin motif (which corresponds to the putative fusion peptide ^28–30^), whilst the C-terminus is composed of the transmembrane domain and proximal helices. These N and C terminal modules are linked via a loop that bridges around and across the top of the protein. The very N-terminus of E1 forms a structurally divergent tail, which may mediate species-specific interactions with E2 (see below). Notably, in this arrangement the helical hairpin is packed alongside the transmembrane domain, close to, or within, the presumed location of the membrane.

**Figure 2.**
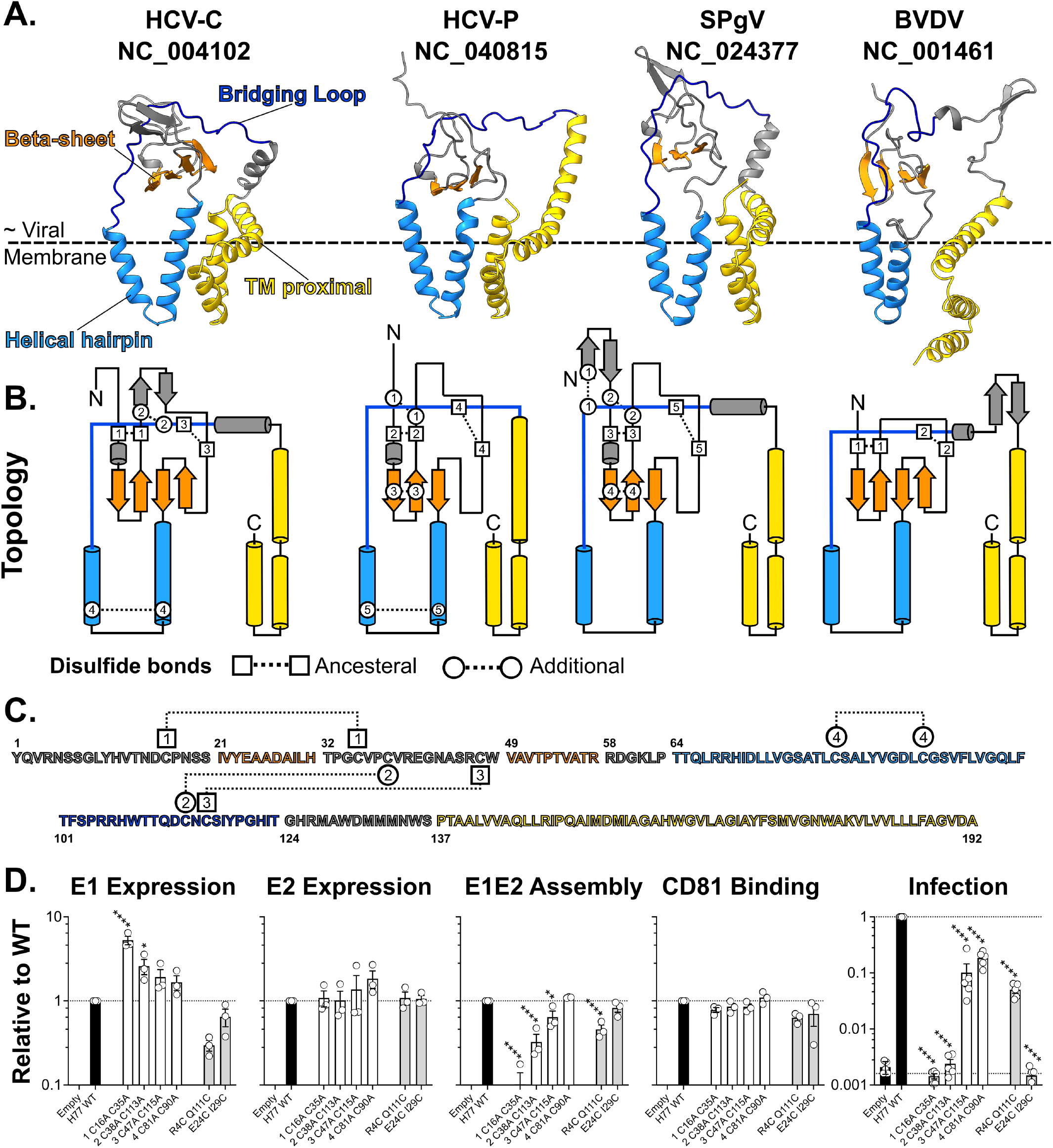
The molecular architecture of E1. **A.** Ribbon diagrams of predicted E1 structures for HCV-C, HCV-P, SPgV and BVDV, features are colour coded as shown on the HCV-C structure, left. The approximate location of the outer leaflet of the viral membrane is inferred by the positions of the E1 transmembrane domain. **B.** Topology diagrams of above. Disulfide bonds are annotated as ancestral (found across all genera) or additional (genera/virus specific). **C.** Linear representation of HCV-C E1 sequence demonstrating the position of each disulfide bond. **D.** HCV E1 was mutated to remove each of its four disulfide bonds, in turn, or to add additional bonds found in Pegiviruses. Each mutant E1E2 was assessed for expression (either E1 or E2), CD81 receptor binding, E1E2 assembly and pseudotyped virus infection. Data is expressed relative to H77 wild type isolate E1E2, n?3. Asterisks indicate degree of statistical significance compared to WT (One-way ANOVA).

E1 is crosslinked by various intramolecular disulfide bonds (Fig. 2B), two of which appear to be ‘ancestral’ being present in each genus. One of these ancestral bonds sits across the inward and outward chains that form the first beta-strand, the other secures the bridging loop to the central beta-sheet. E1 from Pestiviruses possess only these ancestral intramolecular bonds; some Pestiviruses (e.g. Atypical porcine Pestivirus) even lack the second of these bonds. Hepaci and Pegi viruses have various additional bonds, many of these, however, appear to reiterate the interactions imposed by the ancestral bonds; sitting within and proximal to the beta sheet, or linking the bridging loop to the N-terminal region of E1. One additional disulfide bond crosslinks the tip of the helical hairpin motif, this is specific to the Hepaciviruses.

The HCV-C E1 model is in agreement with experimental structures of various E1 peptide fragments and, importantly, the recently reported cryoEM model of E1E2 ^31^; for example disulfide bonds 1, 2 & 3 are resolved, and confirmed, in this structure (Fig. S16A & B). However, our model, and the recent cryoEM model, differs significantly from a previous crystal structure of the N-terminal region of E1 (Fig. S16C); whether this represents a crystallisation artefact or a functionally relevant alternative conformation requires further investigation. Systematic comparison of our predicted E1 structures with entries in the Protein Data Bank and the AlphaFold/EBI Structure Database revealed no significant structural homology. This suggests that E1 exhibits a unique protein fold. Moreover, the lack of similarity between E1 with other viral glycoproteins, is consistent with the notion that the Hepaci, Pegi and Pestiviruses possess a novel fusion mechanism.

We explored the importance of E1 disulfide bonds through mutagenesis and antigenic/functional characterisation in HCV-C E1E2 pseudotyped viruses (Fig. 2C & D). Individual loss of any of the four HCV-C E1 disulfide bonds did not negatively impact expression of E1 and E2, or affect binding to the CD81 receptor (which occurs via E2, without contribution from E1). However, loss of C16-C35, an ancestral bond, or C38-C113, an additional bond, prevented E1E2 dimer assembly and infection. Notably, C47-C115 (ancestral, connecting the bridging loop and beta-sheet, missing in some Pestiviruses) and C81-C90 (additional, crosslinking the helical hairpin) were not absolutely essential for E1E2 dimerisation or infection.

To further explore our models, we mutated HCV-C E1 to introduce new disulfides, analogous to the additional bonds found in E1 from distantly related Pegiviruses (Fig. 2D). R4C-Q111C and E24C I29C (analogous to bonds 1 and 4 in the SPgV E1 model), had limited impacts on E2 expression, CD81 binding and E1E2 dimerisation; this suggest the positioning of these bonds are compatible with the predicted topology and structure of E1. R4C Q111C permitted infection, albeit reduced (potentially due to lower levels of E1), whereas E24C I29C was non-infectious despite normal expression/folding; this may indicate that E1E2 are locked in a properly folded, but inactive, state. An important context to the patterns of folding and activity revealed by these experiments is that disulfide isomerisation may be important for Hepacivirus (and Pestivirus) entry ^32–34^. Therefore, some of these ancestral and additional bonds may be broken and/or shuffled during the conformational rearrangements necessary for fusion. These models, and experiments, provide a platform for future investigations in this area.

## Structural conservation in E2

The high degree of predicted structural homology in E1 would suggest that it performs a mechanistically conserved role in membrane fusion, consistent with it possessing the putative fusion peptide/loop ^28–30^. In contrast, E2 exhibits high structural divergence between, and even within, viral genera. This, most likely, indicates that E2 mediates species-specific interactions with a particular host; indeed, each genera varies widely in tissue tropism (e.g. Hepaciviruses exclusively infect the liver, whilst Pegiviruses target bone marrow). Nonetheless, unbiased structural comparison identified some consistent elements across divergent E2 models (Fig. 3 A & B). Hepaci and Pegivirus E2 share a beta-sandwich fold with conserved topology and structure (originally reported in the experimentally-determined models of HCV-C ^23,24^). This central element appears to act as a scaffold around which various species-specific extensions and loops are arranged. This beta-sandwich is, however, absent from Pestiviruses. Another consistent, yet minimal, feature is an extended beta strand that corresponds to the Back Layer apparent in the crystal structures of HCV-C E2 ^23,35^. This region is important for HCV entry ^35^ and is present in E2 models across each genus, suggesting mechanistic conservation. Finally, in all cases the transmembrane domain is predicted to adopt a helical hairpin conformation.

Notably, we also identified structural similarity between E1 and E2 (Fig. 3C & D). The topology and arrangement of the E1 Beta-sheet and Helical hairpin (the N-terminal module, described in Fig. 2), is mirrored in the C-terminal stem region and transmembrane domain of E2. This is best exemplified by comparison of the E1 and E2 beta-sheet elements across viral species; here, unbiased structural comparison supports similarity between the N-terminal beta-sheet of Pestivirus E1 and the C-terminal beta-sheet of Hepacivirus E2 (Fig. 3E - note areas of similarity in orange - and 3F). We propose that this topological/structural similarity may point to the evolutionary origin of E1E2, with an ancient genetic duplication event from a progenitor E1-like fusion protein giving rise to a proto-E2 molecular partner, analogous to the domain duplication event evident in the N- and C-terminal domains of retroviral capsids ^36,37^. The E1E2 of Hepaci/Pegi/Pestiviruses would have all arisen from this common ancestor. In this scheme, E1 is responsible for the primary fusion activity and, therefore, has a conserved structure, whereas E2 performs a regulatory role and has been free to evolve structural elaborations that permit diversification of host and tissue range.

**Figure 3.**
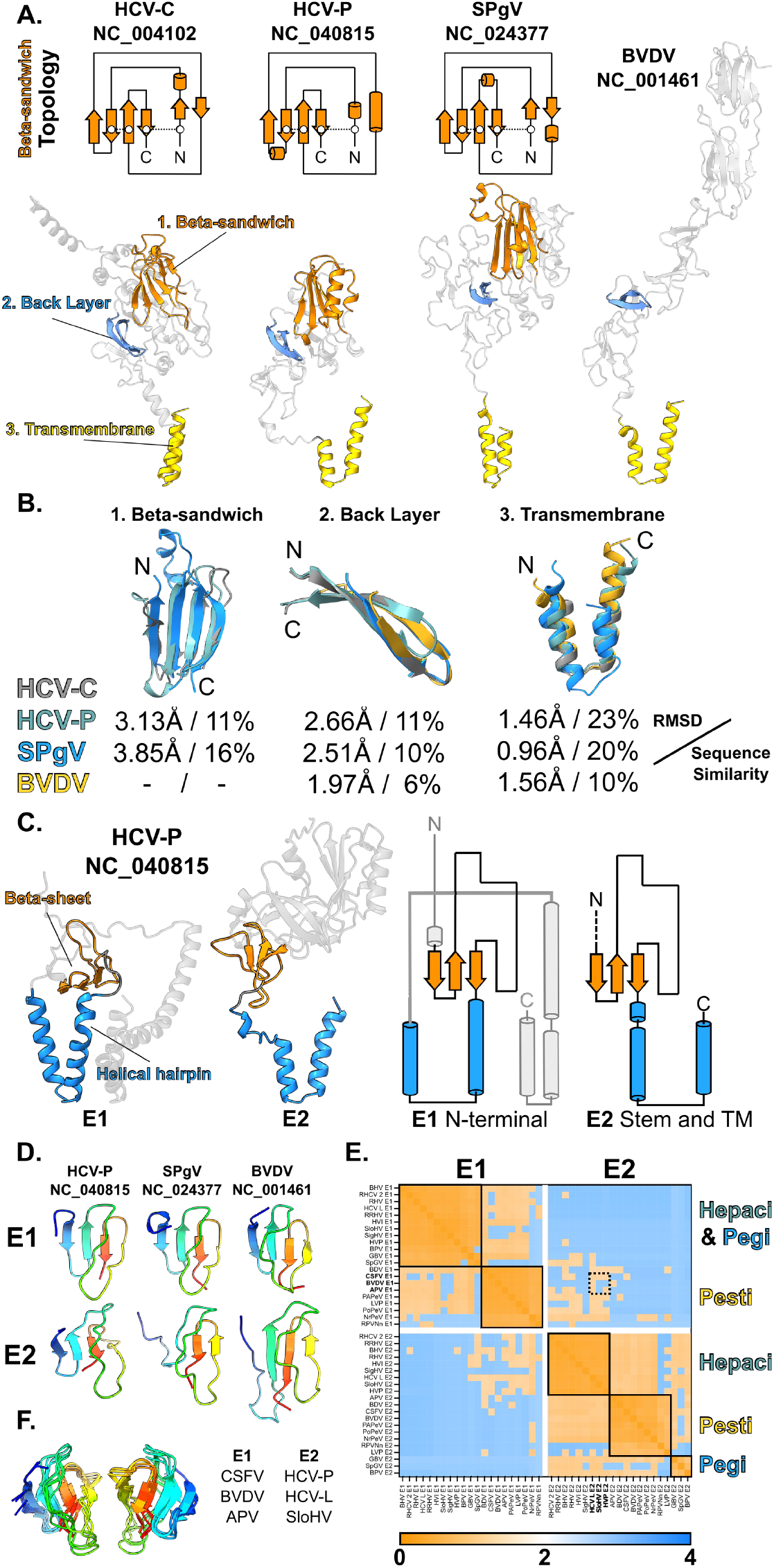
Structural conservation in E2. **A.** Bottom, ribbon diagrams of predicted E2 structures for HCV-C, HCV-P, SPgV and BVDV, consistent features are colour coded as shown on the HCV-C structure, left. Top, topology diagrams for the conserved beta-sandwich scaffold found in all Hepaci and Pegiviruses. **B.** Ribbon diagrams of overlaid features, for HCV-C, HCV-P, SPgV and BVDV, are accompanied by RMSD and sequence similarity values by comparison to HCV-C. **C.** Structural conservation between the N-terminus of E1 and the C-terminal stem and transmembrane domain of E2. Ribbon (left) and topology (right) diagrams of predicted HCV-P E1 and E2 structures highlighting similarity. **D.** Ribbon diagrams (rainbow colour coded blue to red, N to C terminus) of beta-sheet feature conserved across E1 and E2 in HCV-P, SPgV and BVDV. **E.** Structural similarity heatmap for Hepaci, Pegi and Pesti E1 and E2 beta-sheet elements (as shown in D.); orange denotes low distance and, therefore, high similarity. **F.** Structural conservation between Pestivirus E1 and Hepacivirus E2 beta-sheets demonstrated by superposition of stated structures (annotated as dashed box on E.). Structures are shown in two opposite orientations, rainbow colour coding from N to C termini.

## The E1E2 interface

It has long been established that the folding and activity of E1E2 are interdependent and we expect conserved intermolecular communications to regulate and mediate membrane fusion. Our models indicate that E1 wraps tightly around the transmembrane domain and stem regions at the C-terminus of E2 with an interface composed of multiple E1 regions (Fig. 4A & B). For example, the E2 transmembrane domain packs against the helical hairpin of E1 whilst the E2 stem region abuts the bridging loop of E1. These various interactions create an ‘ancestral’ interface found in all E1E2 (Fig. 4C) and are consistent with the ‘stem in hand’ model based on the recent partial structure of HCV-C E1E2 ^31^. Notably, this ancestral interaction is centred around the region of E2 that, we propose, originated through genetic duplication of E1 (Fig. 3C).

**Figure 4.**
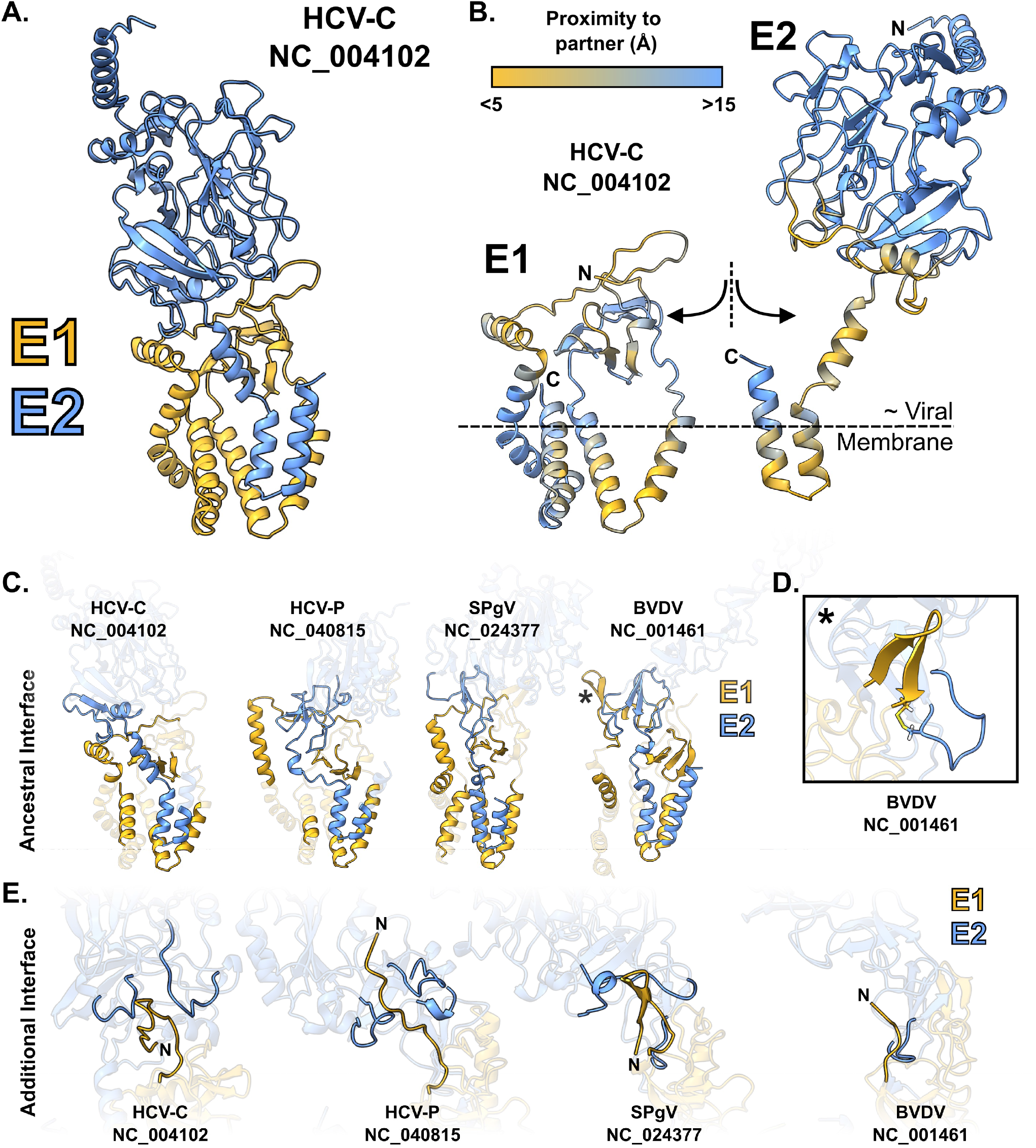
The E1E2 interface. **A.** Ribbon diagram of HCV-C E1E2 complex. **B.** HCV-C E1E2 complex interaction interface. Residues are colour coded by their shortest distance to the partner protein (Cα to Cα). The approximate location of the outer leaflet of the viral membrane is inferred by the positions of the E1 and E2 transmembrane domains. **C.** Ribbon diagrams demonstrating the ‘ancestral’ E1E2 interface found in all genera, regions of contact/high proximity have been highlighted. **D.** Pestiviruses possess an E1E2 intermolecular disulfide bond, the position of which is marked by an asterisk on C. **E.** Ribbon diagrams demonstrating the ‘additional’ species-specific interface between the extended tail of E1, found in Hepaci and Pegiviruses, and loops at the base of E2.

Beyond this ancestral interface there are additional genus/species-specific interactions. First, unique to Pestiviruses, is a predicted E1E2 intermolecular disulfide bond involving a beta-hairpin, found directly upstream of the transmembrane proximal region in E1, and a loop extending from the beta-sheet element in the stem of E2 (Fig 4C & D). Second, the very N-terminus of Hepaci and Pegivirus E1 varies in length and conformation between species. This N-terminal tail makes further contacts with species-specific loops extending from the base of E2; this mode of interaction is largely missing from Pestiviruses as they lack an N-terminal extension in E1 (Fig. 4E).

## Discussion

We have combined systematic curation of viral genome sequences with AlphaFold modelling to perform a structural survey of the E1E2 glycoproteins from 60 diverse viruses and provide models for 24 E1E2 complexes. These analyses revealed that the E1 protein from Hepaci, Pegi, and Pestiviruses exhibits a unique, but very consistent, structure; this suggests that these viruses share a novel viral membrane fusion mechanism.

We have high confidence that these predicted structures are informative: AlphaFold performed very well in benchmarking tests (Fig. S1), our predictions indicate very consistent E1 structures from completely independent datasets (Fig. 1), and our models compare favourably to the current best understanding of E1 and E1E2 based on experimental structures (Fig. S16). There will, undoubtedly, be some inaccuracies in our models. However, insights gained from the depth and breadth of structural comparisons in this study would be difficult to achieve with classical structural biology.

A recurrent theme in our analysis is that of ‘ancestral’ features of E1E2, found in each genus, augmented with ‘additional’ genus or species-specific elaborations. Delineating the contributions made by ‘ancestral’ elements is likely to provide fundamental mechanistic insights. In this respect, Pestivirus E1, with only 1 or 2 intramolecular disulfide bonds and lacking an N-terminal extension, may represent the simplest iteration of this fusion mechanism. This would also be consistent with the ancient ancestral status of Pestiviruses apparent in phylogenetic analyses ^38^.

Ultimately, whilst this work provides the first evidence of a shared novel fusion mechanism in these viral genera, there are many important questions left to answer. Understanding how E1E2 performs membrane fusion will require a detailed mapping of structure to function and an appreciation of the necessary conformational rearrangements. We do not know whether AlphaFold delivers a pre-or post-fusion structure (or some intermediate form). The close packing of the E1 and E2 transmembrane domains with the putative fusion peptide (E1 helical hairpin) may suggest a post-fusion arrangement, but similarly, the agreement of our models with various experimental structures, which required protein stabilisation with neutralising antibodies, supports the notion of a pre-fusion state.

Furthermore, there remain overarching questions around the evolution of membrane fusion in these viruses. Structural/topological similarity between E1 and E2, identified here, may suggest that E1E2 originated in an ancient genetic duplication event; might a primordial fusion apparatus, only containing E1, exist elsewhere in the virome or beyond? Also, how does Hepaci/Pegi/Pestivirus evolutionary history relate to the Flavivirus genus (distant ancestors, also in the Flaviviridae family), which exhibit a completely different class-II membrane fusion protein. Did these genera diverge from one another due to the acquisition of alternative membrane fusion mechanisms?

We expect the structural insights gained through this study will provide a platform for wide-ranging investigations on these questions, and further our understanding of viral membrane mechanisms. This will not only build fundamental knowledge but also guide structure-based design of antiviral interventions and vaccines.

## Data Availability

All AlphaFold structures and associated protein sequences are available here: http://doi.org/10.5281/zenodo.7221315

## Acknowledgements

JG is supported by a Sir Henry Dale Fellowship from the Wellcome Trust and Royal Society (107653/Z/15/Z) and core support from UKRI/Medical Research Council (MC_UU_12014).

## Author Contributions

Conceptualisation: JG. Investigation: MO, KT, CL, SD, RG, JG. Formal Analysis: MO, RG, JG. Data Curation: MO, JG. Visualisation: JG. Writing: JG, KT. Supervision: JG. Funding: JG.

## Notes

http://doi.org/10.5281/zenodo.7221315

### Competing Interest Statement

The authors have declared no competing interest.

