## Supplementary Materials for "Evidence of a novel viral membrane fusion mechanism shared by the Hepaci, Pegi and Pestiviruses"

#### Generation and Assessment of AlphaFold E1E2 Models

##### Background

Despite significant technical challenges around protein production and stabilisation, experimental structural biology has provided an increasingly clear picture of the major glycoprotein, E2, of Hepatitis C Virus (HCV-C) and Bovine Viral Diarrhoea Virus (BVDV) [1–5]. These analyses revealed significant structural divergence in E2 from these prototypical Hepaci and Pestiviruses, and it has remained unclear how their fusion mechanisms may relate to one another, if at all. Information on the minor glycoprotein, E1, is limited [6] and the architecture of the E1E2 complex is only now starting to be revealed by cryoEM [7] (although experimentally-determined E1E2 models are not yet publicly available). Consequently, there remains a poor understanding of the potential mechanisms of membrane fusion by Hepaciviruses, Pestivirus and, the closely related, Pegiviruses.

Recent advances in protein structure prediction, driven by machine learning, has enabled high-throughput *ab initio* protein modelling using primary sequence data alone [8]. The resulting databases contain millions of computationally predicted protein models, many of which are of high accuracy. Proteins from the virome are, however, poorly represented in these databases; for example the HCV-C glycoproteins are not present. Moreover, many viruses (including the Hepaci/Pegi/Pestiviruses) encode polyproteins that are processed by proteases to liberate their constituent proteins. Consequently, unlike eukaryotic open reading frames (ORFs), the coding sequences of these viral proteins are not simply delimited by start and stop codons and, therefore, are poorly suited to automated data mining and structure prediction. Modelling of structures from such viral species requires careful sequence curation and annotation to identify target proteins within the larger ORF.

We set out to apply AlphaFold [9], the current leading structure prediction method, to investigate the E1E2 fusion glycoproteins of Hepaci, Pegi and Pestiviruses. We hoped to resolve the overarching questions: do these viruses share a fusion mechanism and whether this mechanism is novel. We also expected structure comparison would provide insights on the evolution of these proteins within and across viral genera.

#### Benchmarking

To build confidence in this approach we first generated AlphaFold models of analogous viral protein targets for which experimental structures are already available [1,3–5,10–13]. Using the ColabFold platform [14] we modelled NS3, NS5B and E2 from HCV-C and BVDV; in the case of E2 we used multiple sequences each representing different variants used in previous crystallographic studies. We also modelled E, a prototypical class-II fusion protein, from Tick Borne Encephalitis Virus, a classical flavivirus. Importantly, no structural templates from the protein database were used for this or any subsequent predictions presented in our work. The AlphaFold models were largely indistinguishable from their cognate experimentally-determined structures (Fig. S1A); each being superposed with very low root mean square deviation (RMSD). The most divergence occurred around differential positioning of domains with, otherwise, very similar folds, possibly suggesting flexible linker regions (e.g. BVDV E2 structures, Fig. S1A).

The ColabFold implementation of AlphaFold produces five models per target protein, which are scored and ranked by pLDDT prediction confidence, and allows an optional AMBER relaxation stage [15,16] to resolve unnatural bond angles and atomic clashes. We evaluated these outputs using the MolProbity scoring method [17] as a surrogate measure of structure quality (Fig. S1B-D). AMBER relaxation resulted in a significant improvement MolProbity score and, therefore, was adopted for all subsequent modelling. We also observed a strong linear correlation between the pLDDT and MolProbity score; this suggests that high AlphaFold prediction confidence yields high quality structural models. Finally, rank 1 AlphaFold models had significantly lower MolProbity scores (i.e. higher quality) than their experimentally-determined partners. Taken together these data suggest that AlphaFold is capable of producing high-quality, accurate and informative predictions for viral protein targets, including some of the glycoproteins that we are directly investigating (HCV-C/BVDV E2).

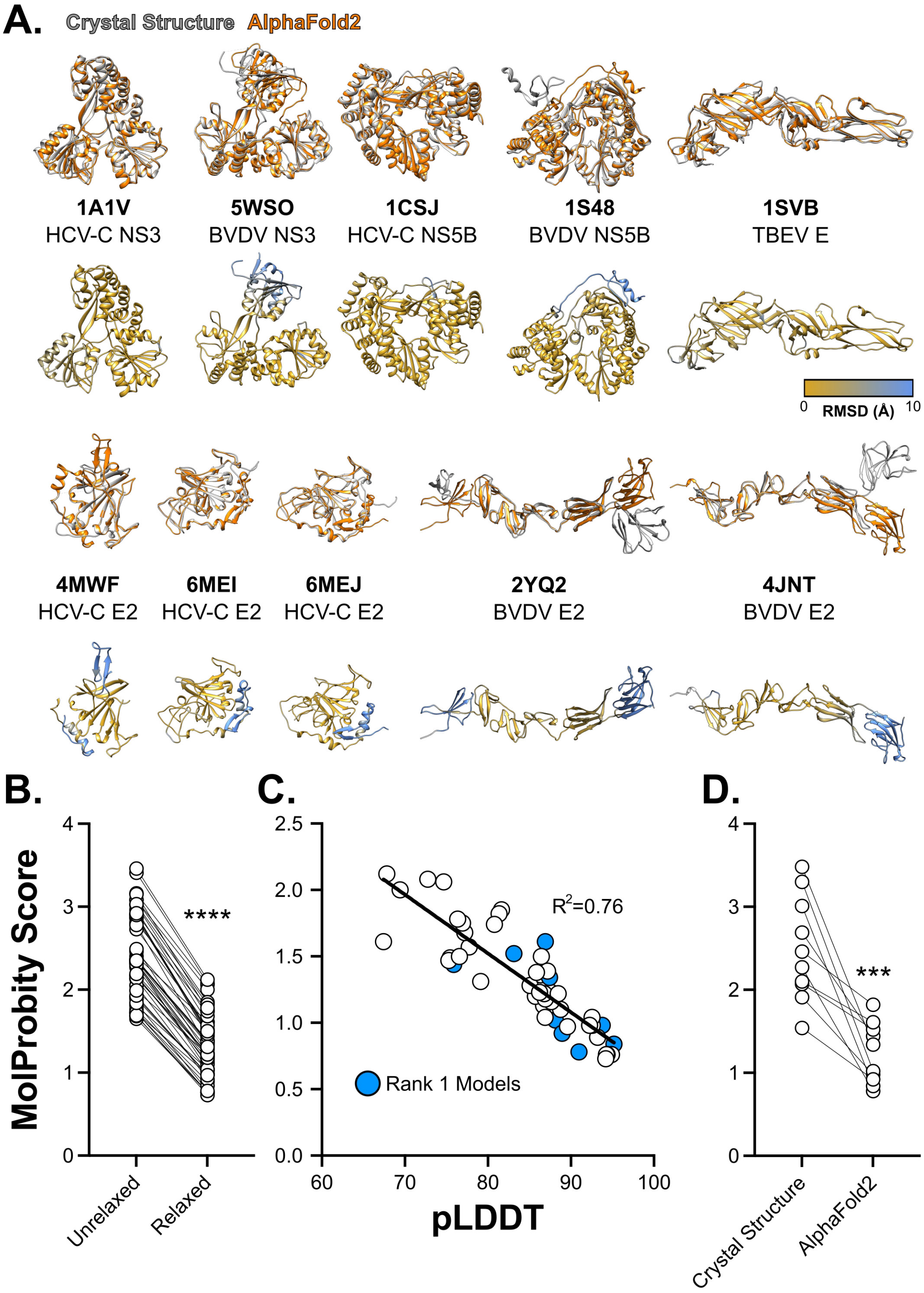


**Figure S1. Benchmarking of AlphaFold against relevant viral targets A.** AlphaFold predicted structures superposed with their cognate experimental structure from the protein database (PDB codes are provided for each structure). In each case the lower structure represents the AlphaFold model colour coded by RMSD from experimentally determined structure. Blue indicates disagreement, as denoted in the colour key. **B.** MolProbity scores of all AlphaFold benchmarking models before and after AMBER relaxation. Each data point represents an individual model, n= 50 (5 candidate models per viral target). Lower values indicate higher model quality **C.** AlphaFold model confidence plotted against MolProbity score for all benchmarking structures (n=50, with rank 1 models shown in blue). The negative correlation indicates higher model quality with higher confidence scores (Linear correlation). **D.** Comparison of MolProbity scores for AlphaFold structures and their cognate experimental structures. Asterisks indicate degree of statistical significance (T-Test).

#### Sequence curation and annotation

High-throughput sequencing of animal and environmental samples continues to identify novel viruses from across the virome. Consequently there are dozens of species from each of the Hepaci, Pegi and Pestiviruses that are available for structure prediction [18]. However, as described above, the N and C-termini of E1 and E2 are delimited by proteolytic cleavage from a polypeptide (which is encoded by a single ORF) and, therefore, individual protein targets need to be defined through sequence curation. For prototypical viruses (e.g. HCV-C, BVDV), pre-existing genome annotations for E1 and E2 are accurate, however, many more-recently discovered viruses lack annotation (or have inaccurate annotations).

To derive unambiguous E1 and E2 sequences for structure prediction we assembled genus-specific alignments of viral genomes, with sequences being drawn from GenBank. This revealed low homology across most of the genome in every genus, thus preventing attempts to propagate protein annotations from prototypical viruses to other species. We therefore reconstructed phylogenies using MSAs spanning only the highly conserved NS5B region. Phylogenies disclosed robustly supported subclades within each genus, we assigned arbitrary names to these subclades and created multiple sequence alignments (MSA) containing just the members of a given subclade. The higher level of similarity within each subclade allowed us to align most of the viral genome including, crucially, the regions encoding E1 and E2. As an example, we provide the Hepacivirus NS5B phylogeny divided into nine subclades (Fig. S2).

We selected master sequences within each subclade (e.g. HCV-C for Hepacivirus clade 1, Fig. S2). For each of these we used the signalP server [19] to predict the signal peptidase cleavage sites that define the termini of E1 and E2, and, where available, cross-referenced predicted sites with pre-existing genome annotations. These locations were then propagated throughout each subclade alignment to generate E1 and E2 sequences for each virus. Finally, we filtered these sequences to remove duplicate viruses (some have two GenBank entries under different names), those with incomplete sequence data across the regions of interest and those that contain ambiguous amino acid residues due to uncertainty in the underlying genome sequence data. This resulted in 32, 15 and 13 complete E1E2 sequences from the Hepaci, Pegi and Pestiviruses, respectively (a complete list of viral species is provided in the underlying data files: http://doi.org/10.5281/zenodo.7221315).

Initial analyses of the E1 and E2 protein sequences foreshadowed their respective structural conservation and divergence apparent in the subsequent AlphaFold modelling. E1 exhibited a consistent length of ~190 residues in all viral species, irrespective of genus. In contrast, the length of E2 varied between and, in the case of the Hepaciviruses, within genera (Fig. S3). For example the E2 of Wenling shark Hepacivirus, presumably an ancient species, is only 214 residues long, whereas E2 from HCV-C is 363 residues long (possessing significant N-terminal extensions compared to many other Hepaciviruses). These observations of protein length, alone, support the notion of E1 as a conserved component, likely constrained by mechanistic importance; whilst E2 may be under fewer mechanistic constraints and is free to evolve host-specific adaptations.

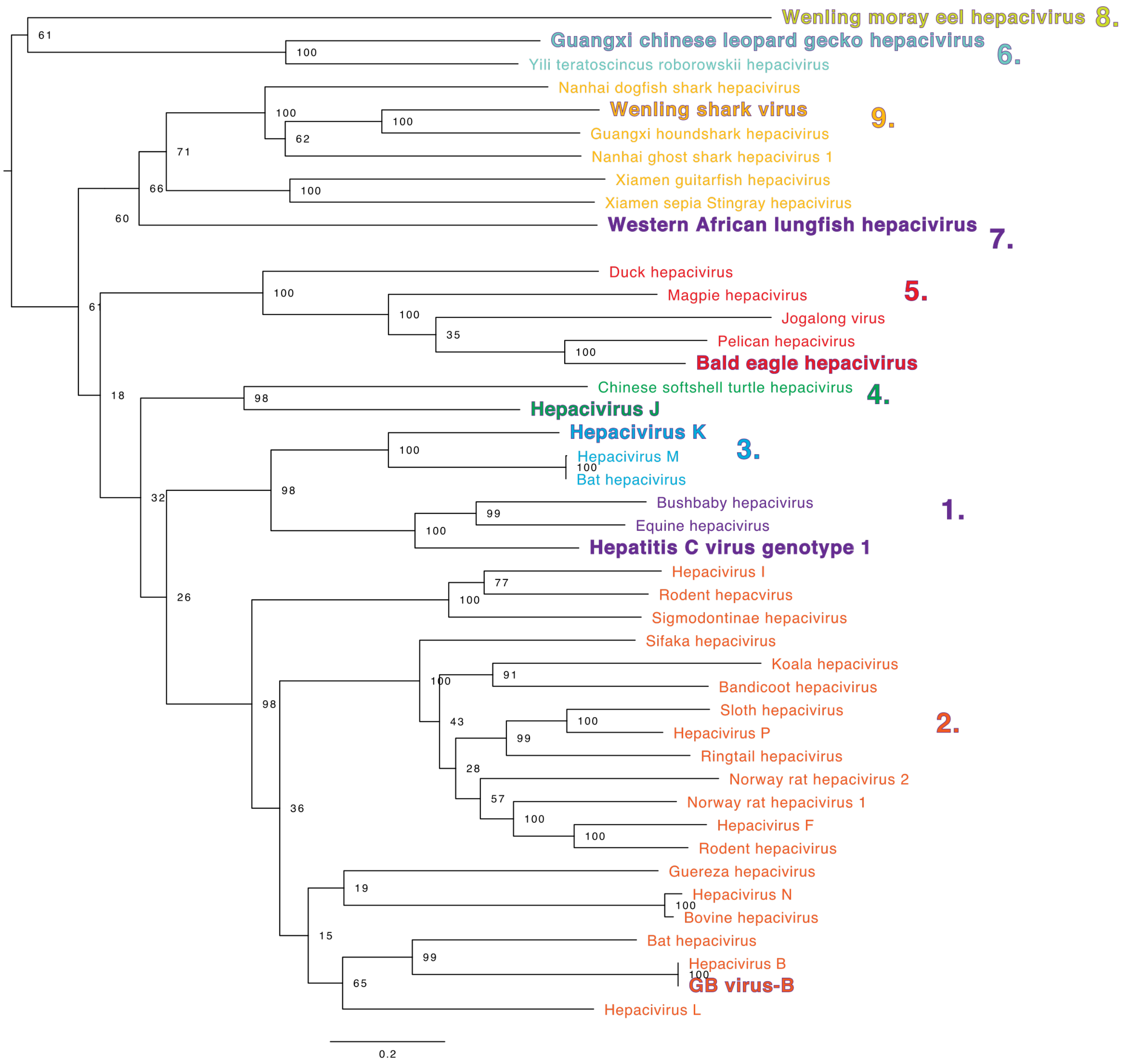


**Figure S2. Hepacivirus NS5B phylogenetic tree.** Subclades are numbered and colour coded; species in bold text represent clade-specific reference viruses. E1 and E2 annotations from these reference viruses were propagated throughout aligned whole genome sequences of each subclade.

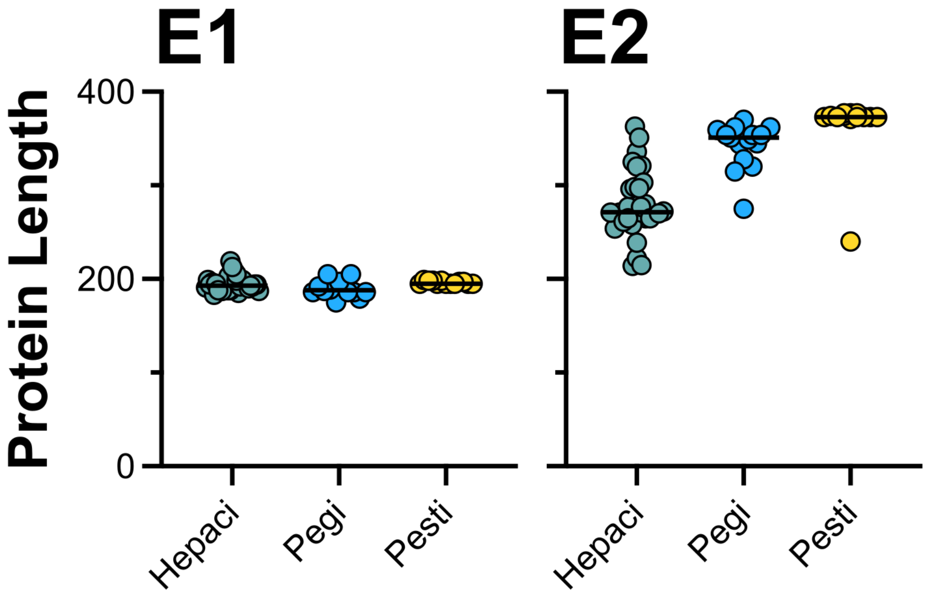


**Figure S3. Protein lengths of E1 and E2 from diverse Hepaci, Pegi and Pestiviruses.** Each data point represents an individual viral species, n=32, 15 and 13 respectively.

#### AlphaFold modelling

In our initial round of modelling we predicted E1 and E2 monomeric structures for all curated viral sequences, we used AMBER relaxation to improve model quality (as outlined in Fig. S1B), and confined our analyses to the rank 1 model for each species (based on pLDDT confidence score). Monomeric E1 and E2 structures for each genus are provided in Figures S4-S9. We evaluated the models using pLDDT and MolProbity scores (Fig. S10). For the Hepaci and Pegiviruses E1 models consistently outperformed their E2 partners in respect to both pLDDT confidence and MolProbity values. Moreover, E2 models displayed a wide range of prediction confidences. This may indicate that the diversity in both sequence and length of E2 erodes the performance of AlphaFold. A simple visual examination of the Hepaci and Pegivirus E1 structures (Fig. S3 and 4) demonstrates a very high degree of structural conservation, which is only lost for very low confidence models. For Pestiviruses, E2 monomer models slightly outperformed E1, this may reflect the high structural uniformity of Pestivirus E2 (Fig. S9). Moreover, whilst E1 structural motifs, conserved in Hepaci and Pegiviruses, are found in Pestivirus E1 models, their overall arrangements differ, with E1 frequently adopting an extended conformation (Fig. S6). Whether these represent mechanistically important confomers requires further investigation.

Guided by this initial round of modelling, we next used AlphaFold to predict E1E2 complex structures. However, given that many E2 monomer models achieved low confidence scores we limited our predictions to those species that scored >70 pLDDT for E2 monomeric structures. This resulted in 11, 3 and, 10 Hepaci, Pegi and Pestivirus E1E2 models respectively (Fig. S11-S13). Model quality metrics suggest that AlphaFold performs somewhat less-well when predicting E1E2 complexes (for example there is a significant reduction in pLDDT scores, Fig. S14). Notably, the conformational heterogeneity apparent in Pestivirus monomeric E1 models (Fig. S6) is eliminated in E1E2 complex models, such that Pestivirus E1 much more closely resembles E1 from Hepaci and Pegiviruses (as presented in the main text Figures 1, 2 and 4). This is consistent with E2 chaperoning the folding of E1 [20].

Finally, we performed unbiased analysis of E1 and E2 structures (from E1E2 complex models) using the all-against-all structural comparison tool on the DALI server [21]. The resultant correspondence analysis from this is provided in main text Figure 1B, whereas Figure S15 displays the dendogram and structural similarity heat map outputs. Here, Hepaci and Pegivirus E1 models group together, with Pestiviruses forming a second group. There is, however, structural similarity between the groups, as demonstrated by intermediate distance values (white and orange) in the heat map. This is consistent with structural conservation in E1. In contrast, E2 structures were divided into four groups: the Hepaci, Pegi and Pestiviruses E2 models and, notably, the HCV-C E2 model alone. The Hepaci and Pegi virus groups, again, share some similarity, but otherwise the groups are quite distinct (demonstrated by higher distance values, coloured in blue). This is indicative of structural divergence. The fact that HCV-C E2 is an outlier, may reflect its evolutionary distance from other Hepaciviruses. Similarly, it may be due to inaccuracies and artefacts in the AlphaFold modelling, although it should be noted that AlphaFold performed well against HCV-C E2 benchmarking targets (Fig. S1). All-against-all analyses on the Dali server was also used to compare E1 and E2 for main text Figure 3E.

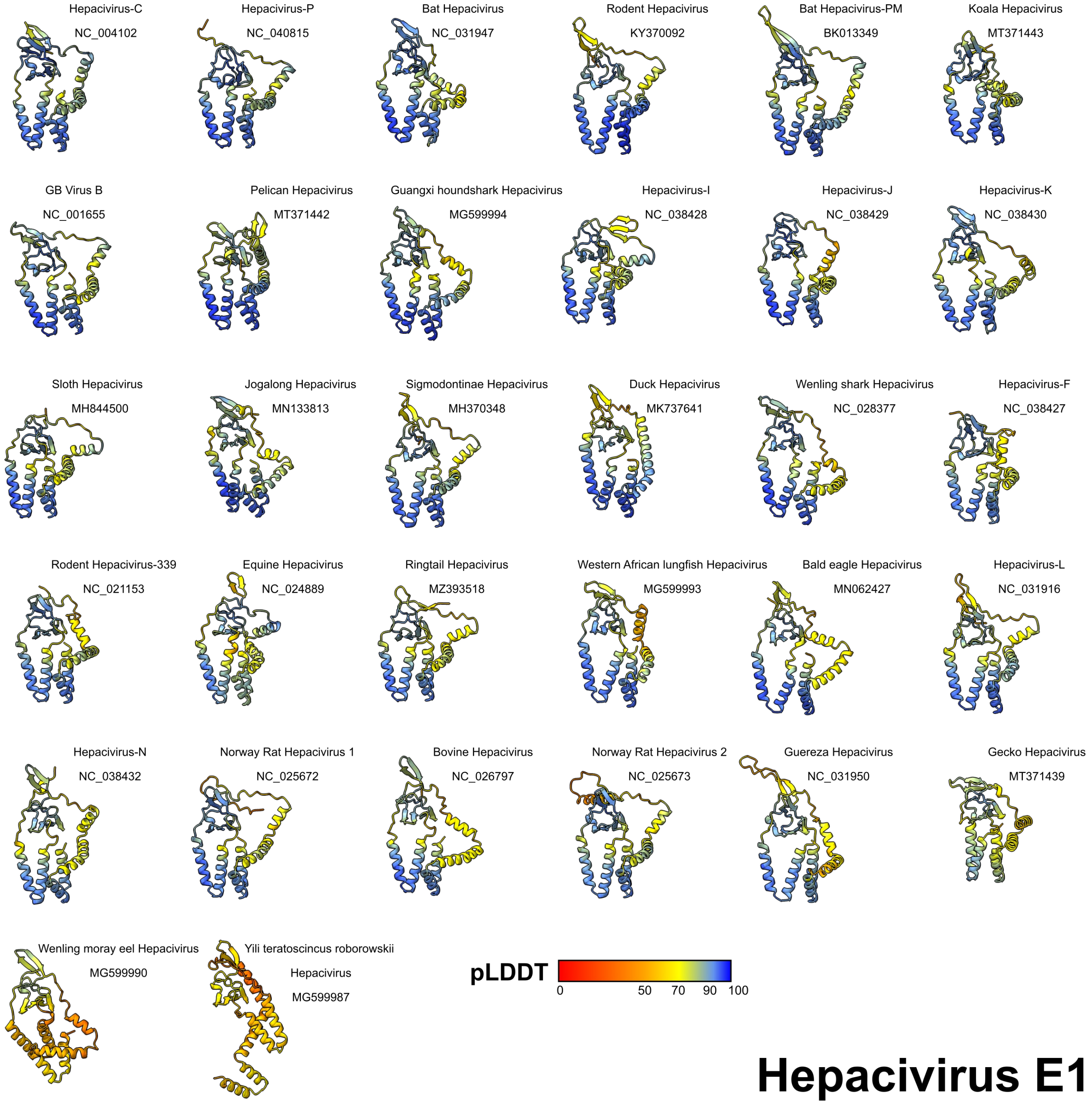


**Figure S4. Hepacivirus E1 monomer AlphaFold models.** Residues are colour coded by pLDDT prediction confidence. Models are arranged in order of descending prediction confidence.

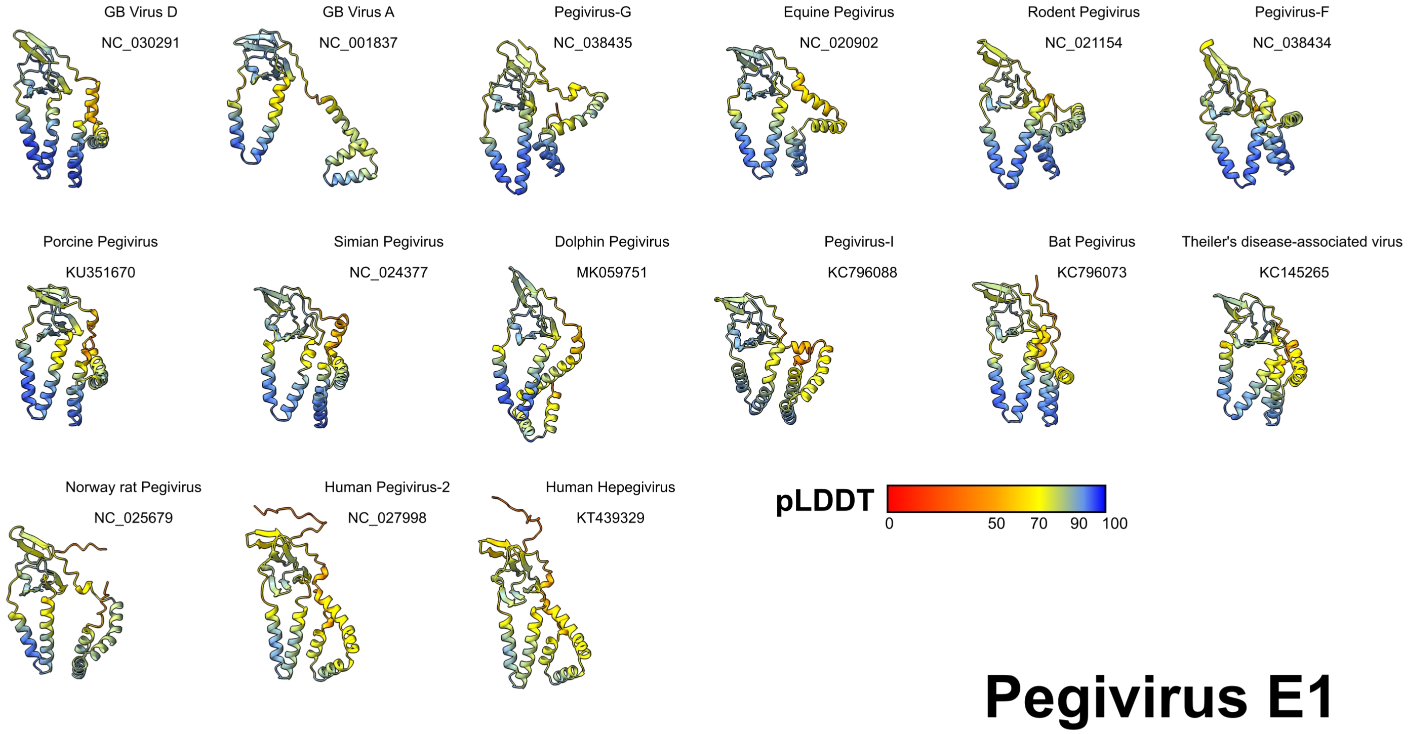


**Figure S5. Pegivirus E1 monomer AlphaFold models.** Residues are colour coded by pLDDT prediction confidence. Models are arranged in order of descending prediction confidence.

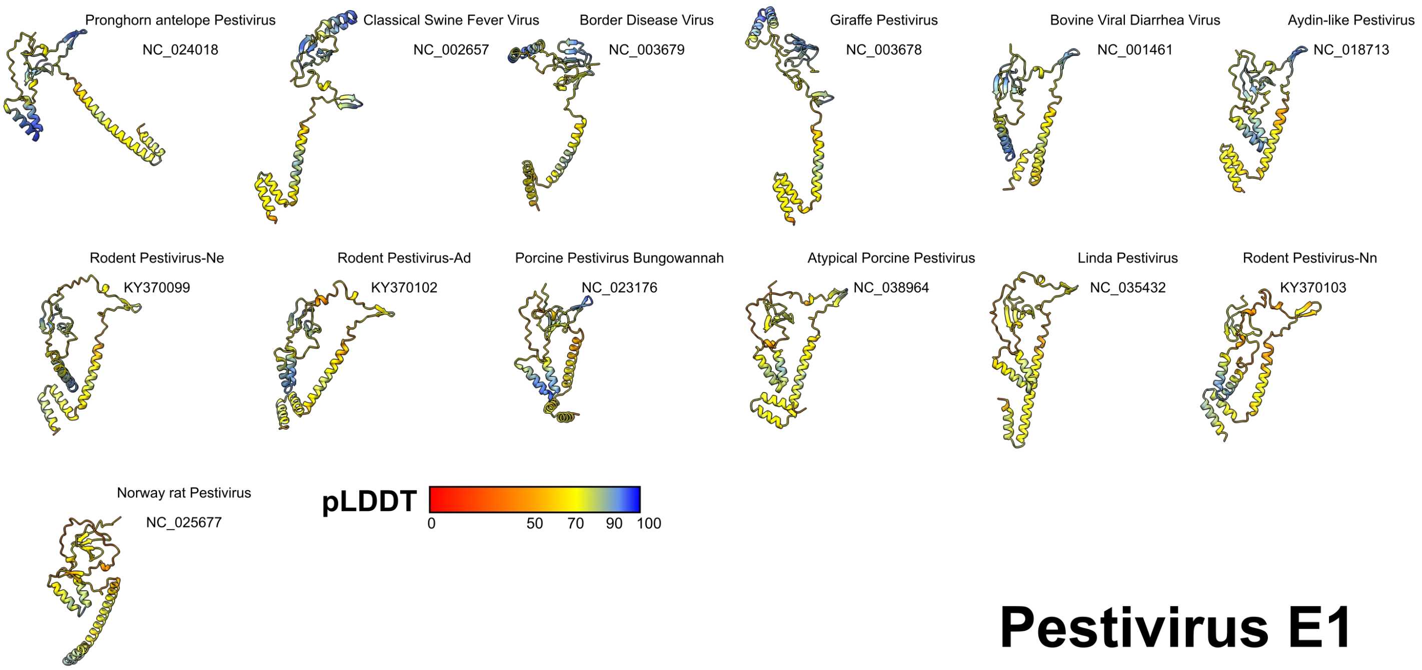


**Figure S6. Pestivirus E1 monomer AlphaFold models.** Residues are colour coded by pLDDT prediction confidence. Models are arranged in order of descending prediction confidence.

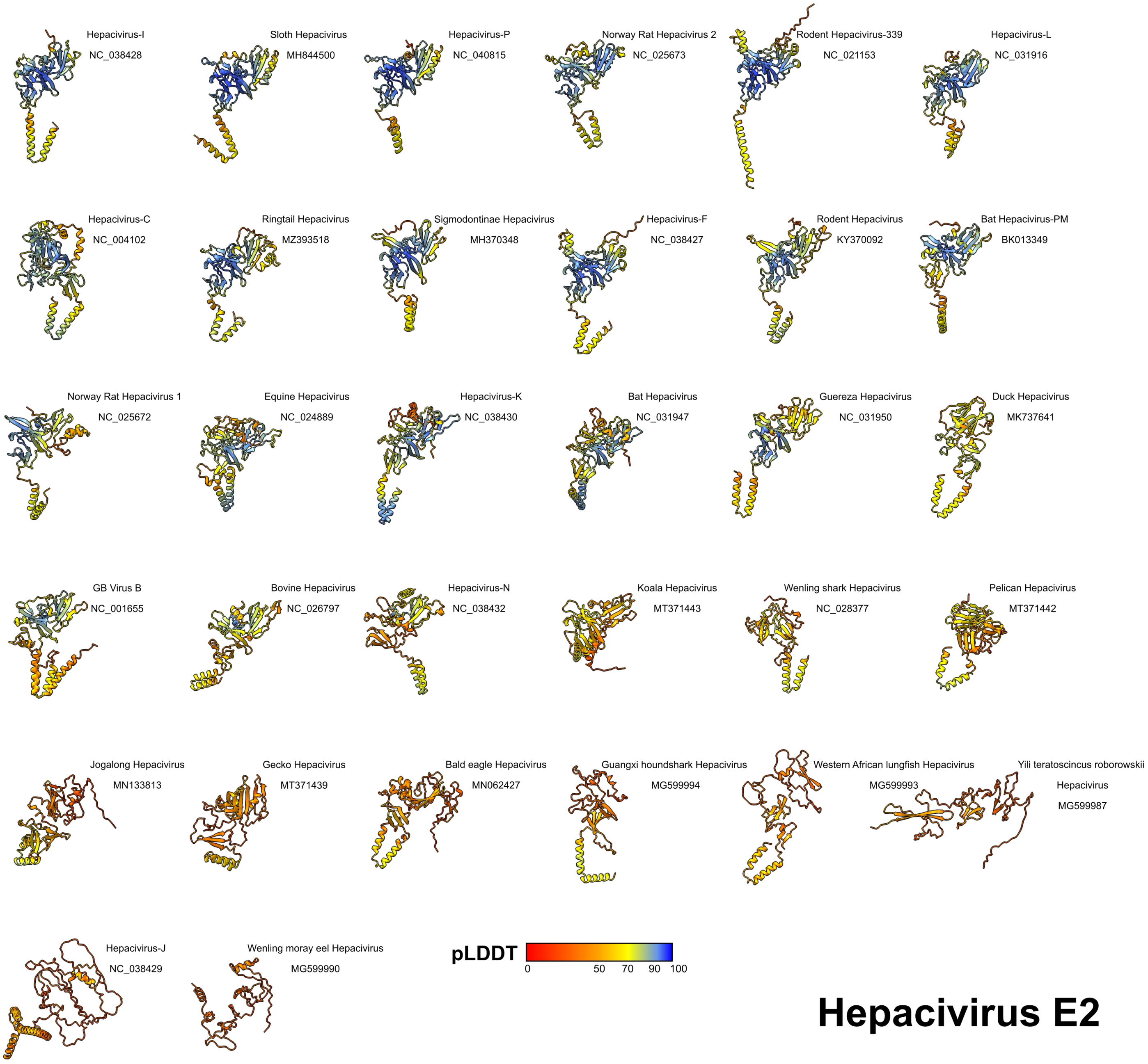


**Figure S7. Hepacivirus E2 monomer AlphaFold models.** Residues are colour coded by pLDDT prediction confidence. Models are arranged in order of descending prediction confidence.

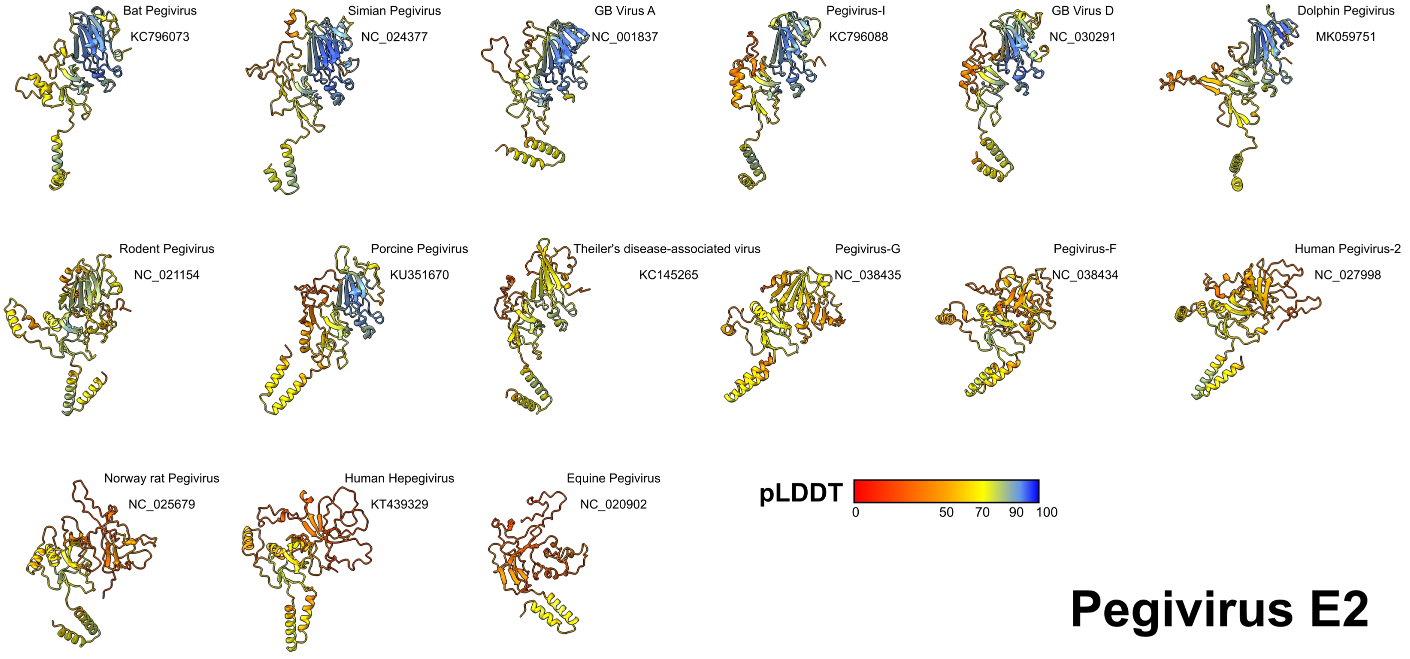


**Figure S8. Pegivirus E2 monomer AlphaFold models.** Residues are colour coded by pLDDT prediction confidence. Models are arranged in order of descending prediction confidence.

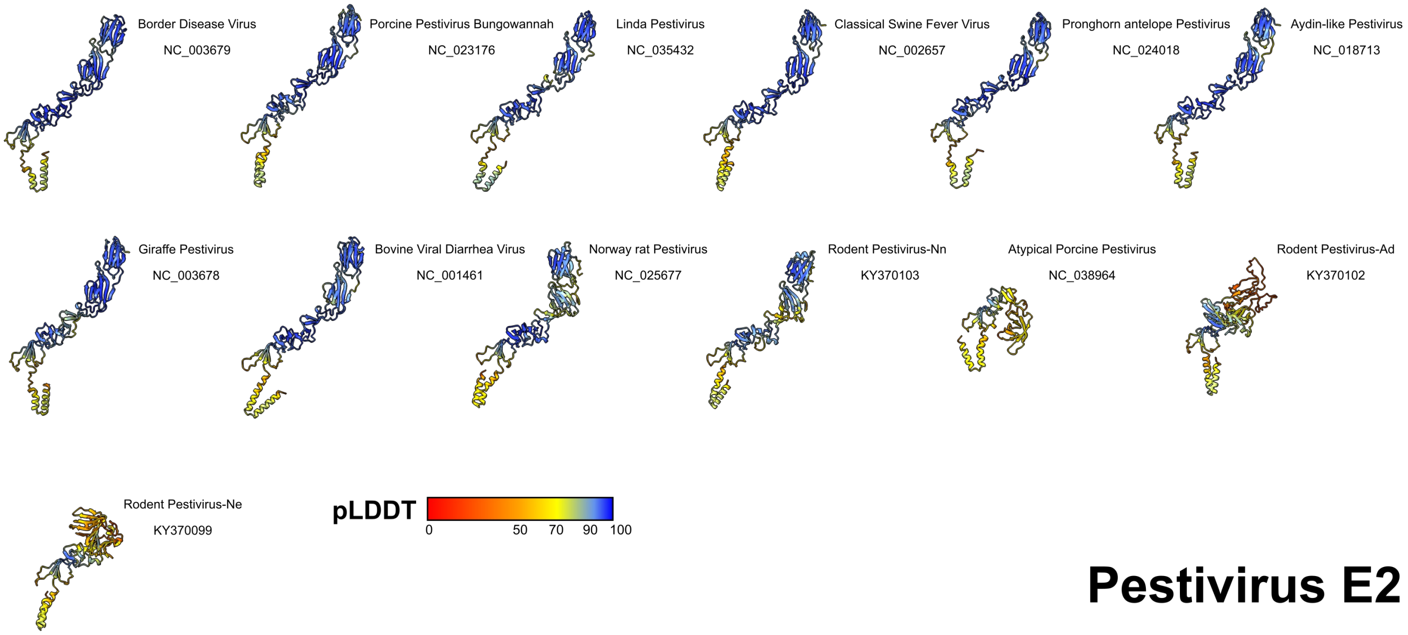


**Figure S9. Pestivirus E2 monomer AlphaFold models.** Residues are colour coded by pLDDT prediction confidence. Models are arranged in order of descending prediction confidence.

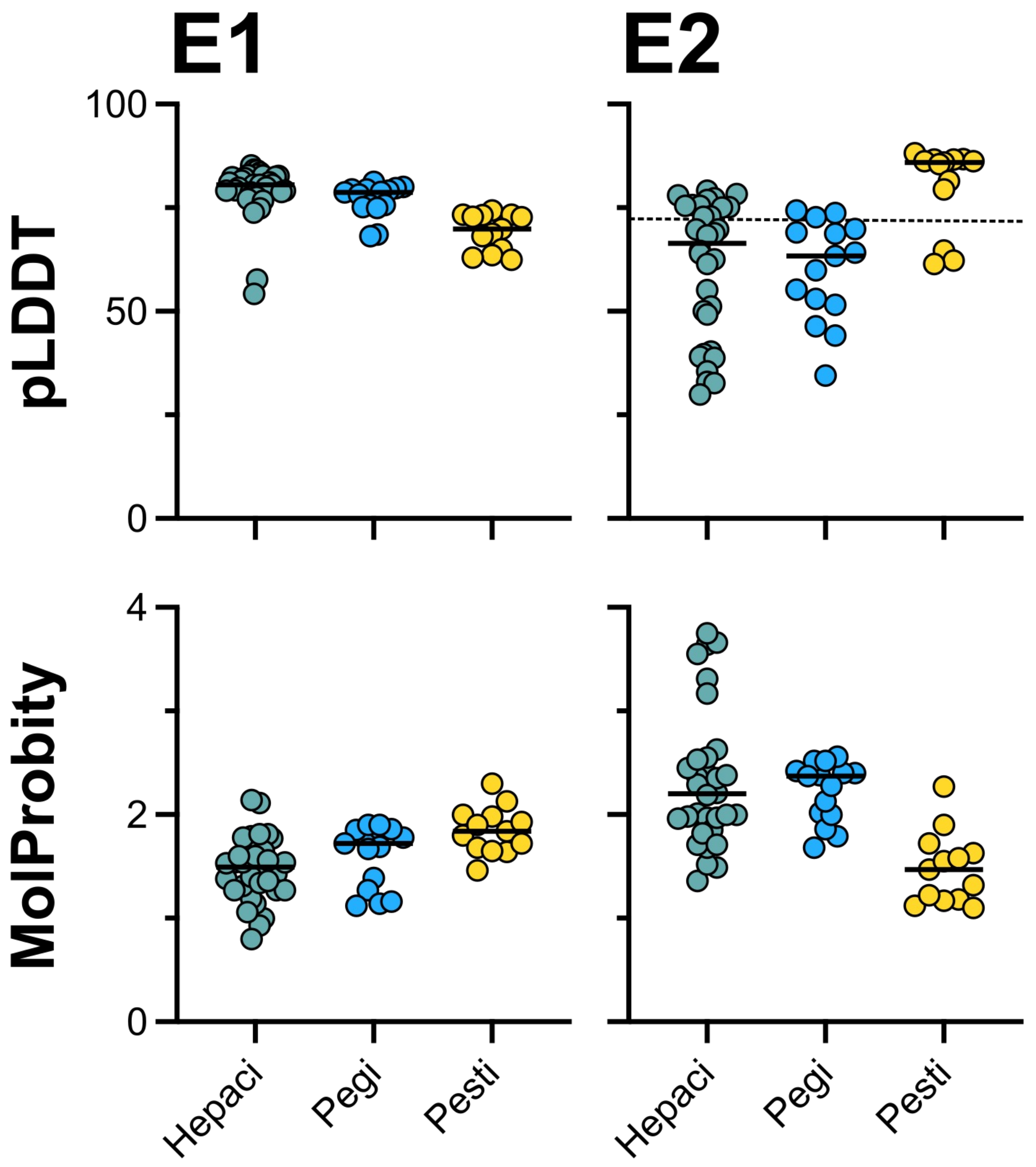


**Figure S10. Model quality metrics for E1 and E2 monomer models.** pLDDT prediction confidence (upper plots) and MolProbity scores (lower plots) for E1 and E2 models from diverse Hepaci, Pegi and Pestiviruses. Each data point represents an individual viral species, n=32, 15 and 13 respectively. Dashed line on upper right plot indicates pLDDT=70 cutoff that was used to select viruses for modelling of E1E2 complexes.

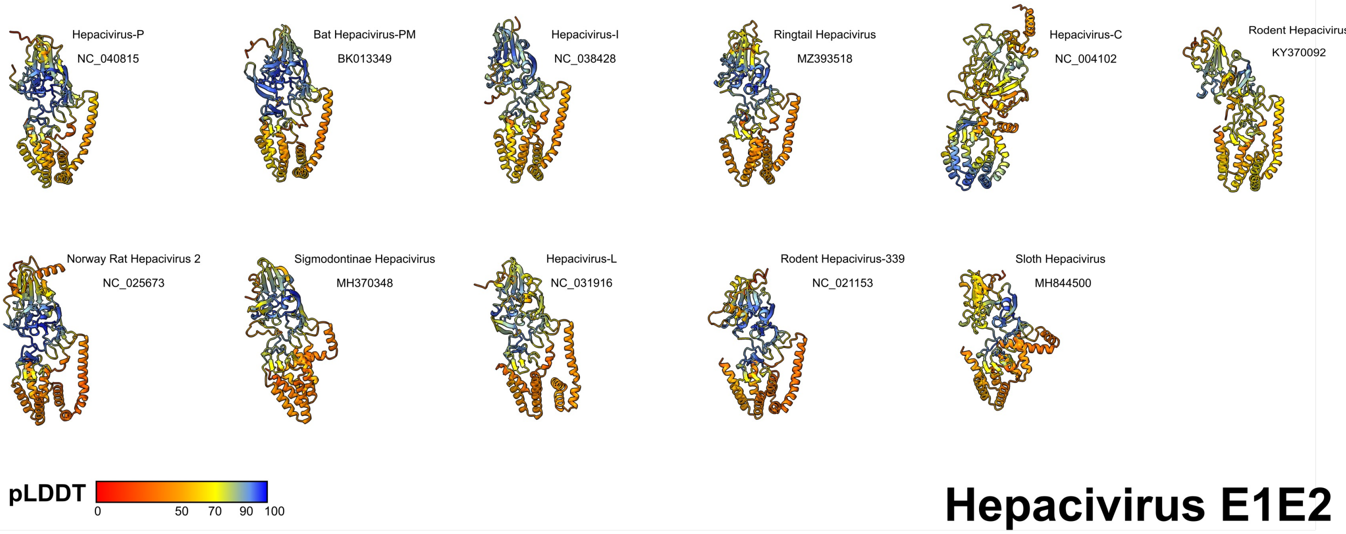


**Figure S11. Hepacivirus E1E2 complex AlphaFold models.** Residues are colour coded by pLDDT prediction confidence. Models are arranged in order of descending prediction confidence.

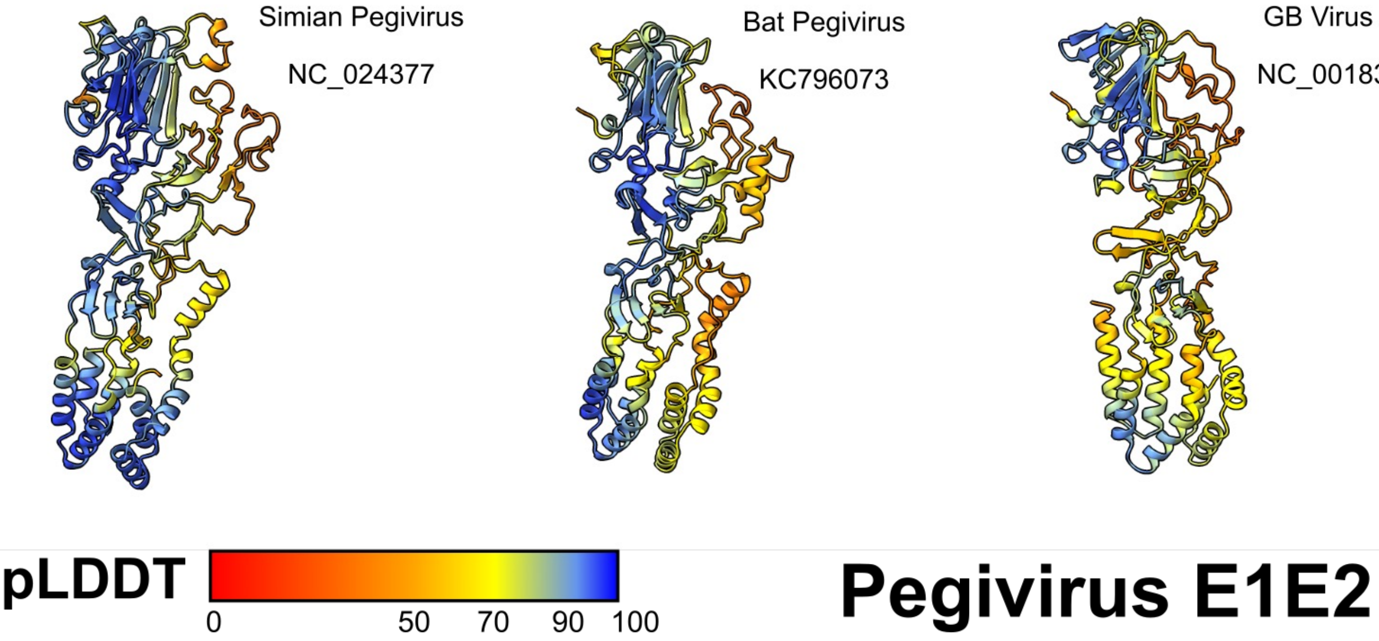


**Figure S12. Pegivirus E1E2 complex AlphaFold models.** Residues are colour coded by pLDDT prediction confidence. Models are arranged in order of descending prediction confidence.

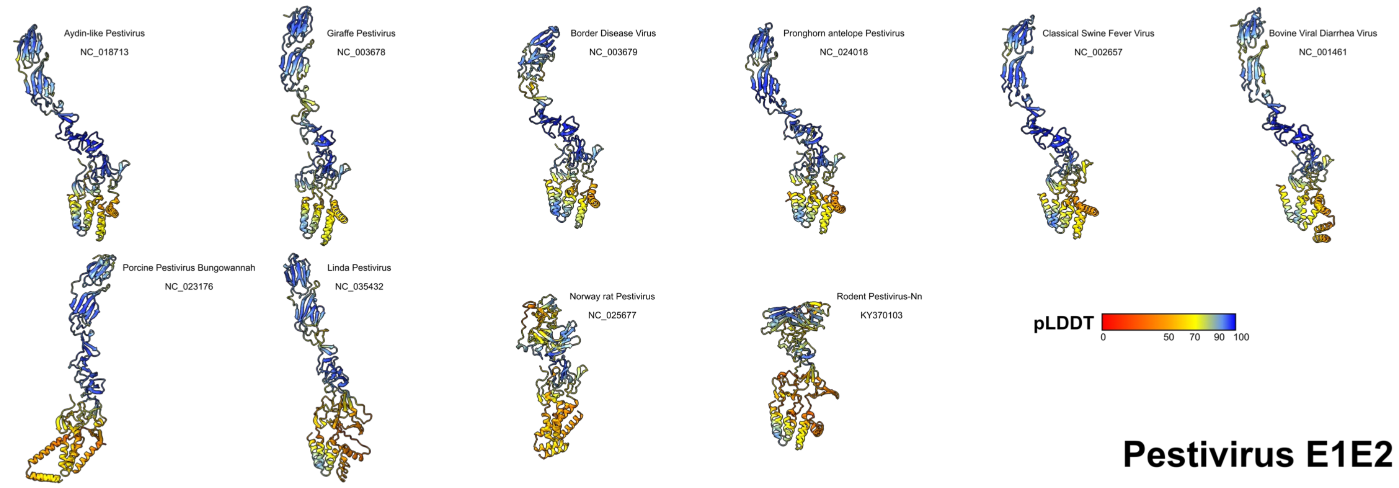


**Figure S13. Pestivirus E1E2 complex AlphaFold models.** Residues are colour coded by pLDDT prediction confidence. Models are arranged in order of descending prediction confidence.

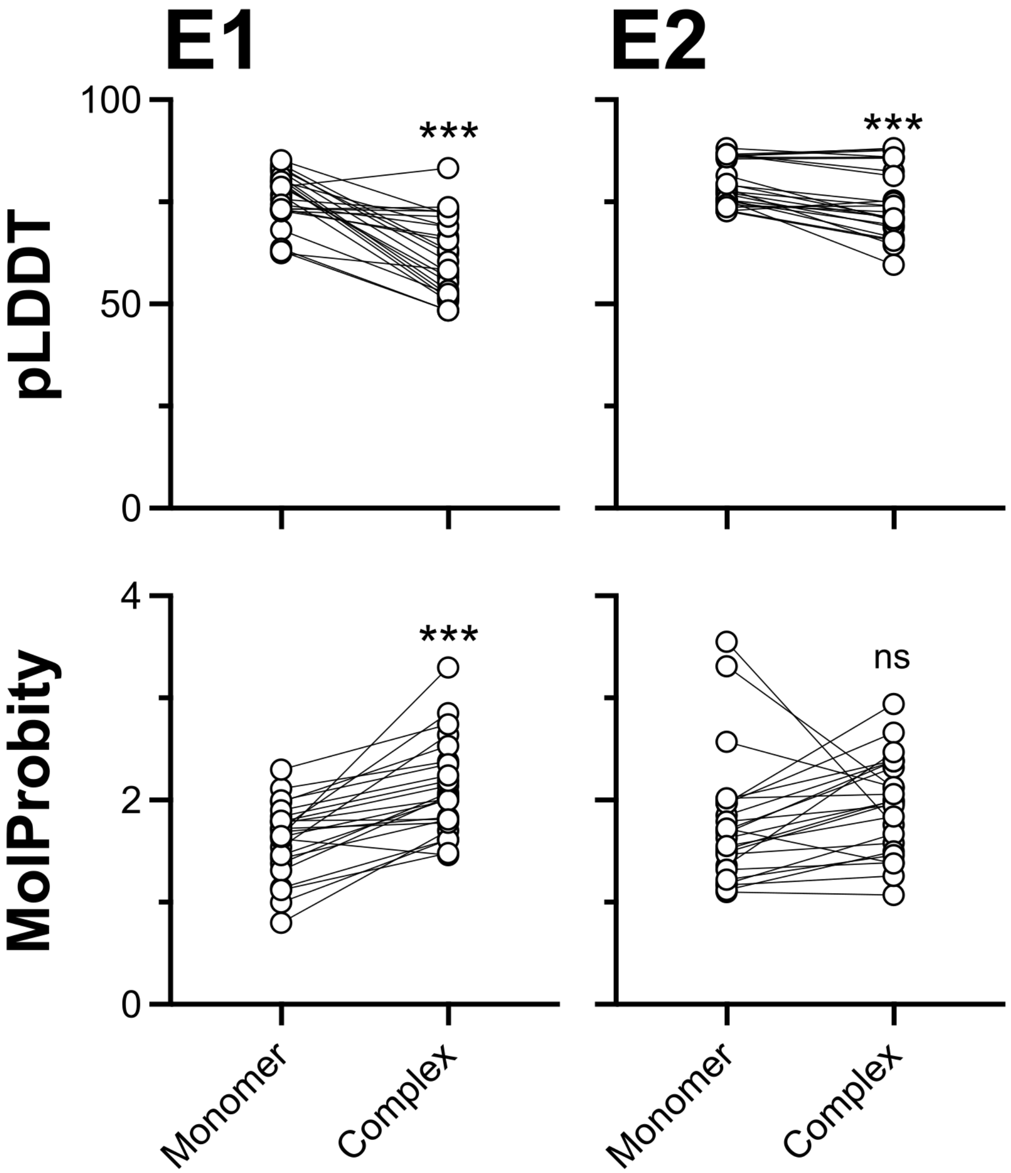


**Figure S14. Model quality metrics for E1 and E2 monomer models.** pLDDT prediction confidence (upper plots) and MolProbity scores (lower plots) for E1 and E2 modelled as a monomer or in a complex. Each data point represents an individual viral species, n=24. Asterisks indicate degree of statistical significance (T-Test).

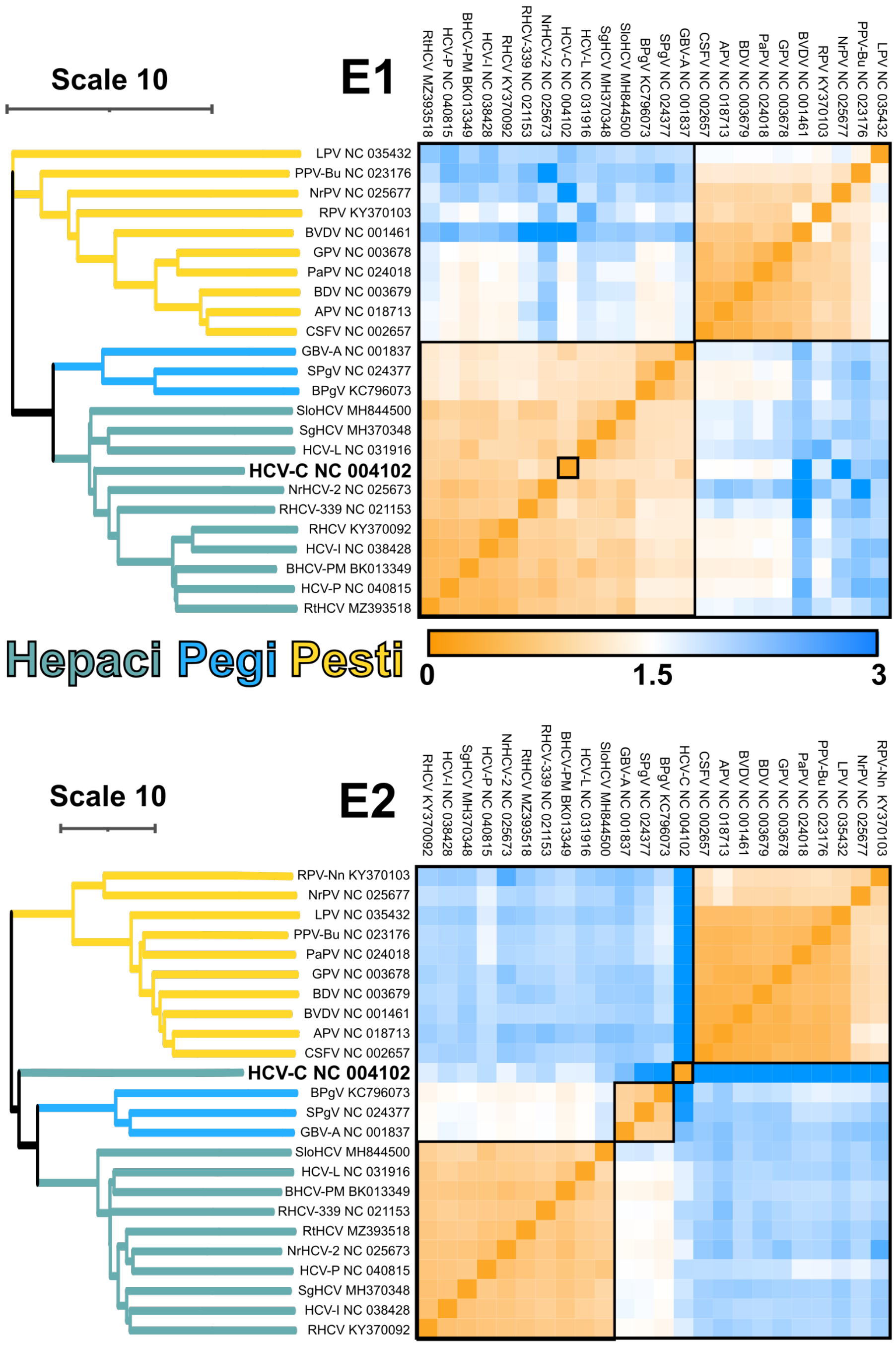


**Figure S15. Unbiased structural comparison and clustering of E1 and E2 models.** All against all comparison was performed on either E1 or E2 (derived from E1E2 complex models) using the DALI server. The heat map (right) provides pairwise distances indicating structural similarity (orange denoting low distances and, therefore, high similarity). Structures are also clustered by similarity, as denoted by the dendogram (left). Boxes are drawn to group structures with high similarity. The position of HCV-C is additionally highlighted.

#### Comparison of HCV-C E1E2 model to experimental structures

We demonstrate in Figure S1 that AlphaFold performs well when predicting the structure of monomeric HCV-C E2. However, there is more limited structural information available for comparison with HCV-C E1 (or E1 from any of the viral species considered here). Nonetheless, NMR spectroscopy and X-ray crystallography of E1 peptides provide some means of model validation (Fig. S16A). Here, three peptides, all focussed on the transmembrane and proximal regions of E1 [22–24], are in very good agreement with with our AlphaFold E1 model (taken from the E1E2 complex structure). The recent partial HCV-C E1E2 cryoEM model, described in a pre-print by Torrents de la Peña and Sliepen et al. [7], would provide an ideal comparator for our predictions but, at the time of writing, this model is not yet available. We can, however, cross reference our model with the disulfide bonding pattern reported in this work, and in previous E2 crystallographic studies [1,3] (Fig. S16B). Here, three out of four predicted E1 disulfides are confirmed in the cryoEM model, with the fourth not being resolved by cryoEM and, therefore, remaining ambiguous. This suggests the topology of our HCV-C AlphaFold E1 model is consistent with experimental data. Moreover, seven of nine predicted E2 disulfide bonds are confirmed in the E1E2 cryoEM model (and at least one crystal structure). Notably, the disagreement in E2 bonding patterns occurs at four closely juxtaposed cysteine residues that have previously been reported to vary in their bonding network (compare 4MWF and 6MEJ, Fig. S16B). This may represent natural and/or functionally relevant diversity in disulfide bonding in this region.

Finally, there is a crystal structure for the N-terminal portion of E1 [6]. Apart from a similar hairpin motif at the very N-terminus, this differs significantly from both our AlphaFold model (Fig. 16C) and, based on disulfide bonding patterns, the recent cryoEM model of E1E2. Given the high consistency of our E1E2 structure predictions across viral genera and their favourable comparisons to other E1 and E2 experimental models, we do not regard that this disagreement invalidates our structural predictions. It may, however, indicate functionally important conformational heterogeneity in the N-terminal region of E1, and will require further investigation.

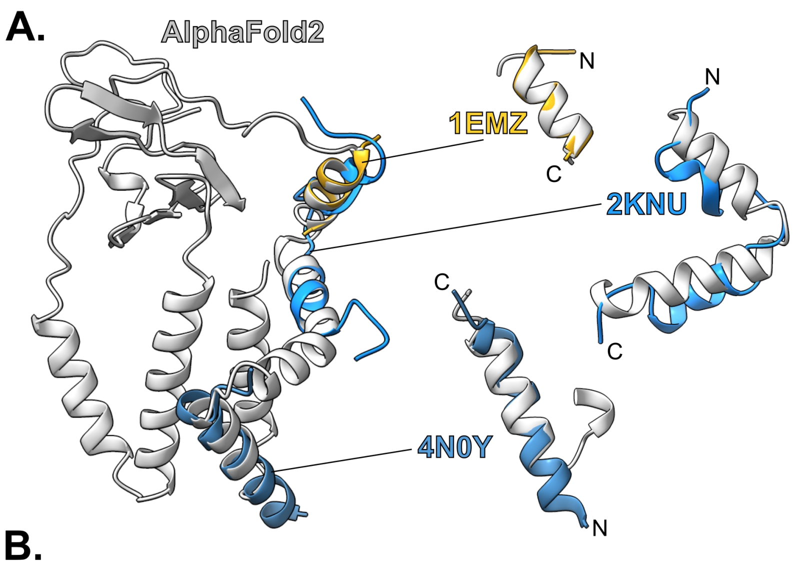


**B.**

| | AF2 E1E2 | CryoEM E1E2 Torrents de la Peña and Sliepen et al. 2021 | 4MWF E2 Kong et al. 2014 | 6MEJ E2 Flyak et al. 2018 |
|---|---|---|---|---|
| **E1** | C207-C226 | C207-C226 | - | - |
| | C229-C304 | C229-C304 | - | - |
| | C238-C306 | C238-C306 | - | - |
| | C272-C281 | NR | - | - |
| **E2** | C429-C503 | C429-C503 | C429-C503 | C429-C503 |
| | C452-C620 | C452-C486 | C452-C486 | C452-C486 |
| | C459-C486 | C459-C486 | - | C459-C486 |
| | C494-C564 | C494-C564 | C494-C564 | C494-C564 |
| | C508-C552 | C508-C552 | C508-C552 | C508-C552 |
| | C569-C581 | C569-C597 | C569-C581 | C569-C597 |
| | C585-C597 | C585-C581 | C585-C597 | C585-C581 |
| | C607-C645 | C607-C645 | C607-C645 | C607-C645 |
| | C652-C677 | C652-C677 | - | - |

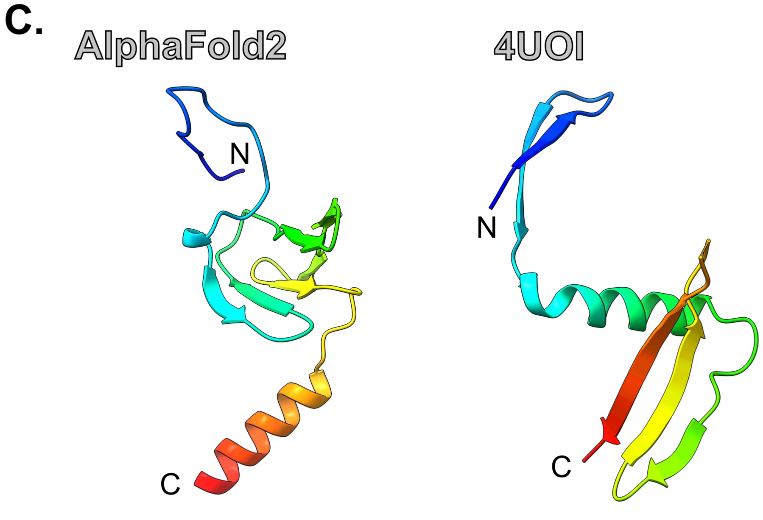


**Figure S16. Comparison of HCV-C AlphaFold E1 model with experimentally determined structures. A.** E1 model compared to cognate peptide structures; superposition was achieved by alignment of structure and sequence. **B.** Comparison of predicted E1E2 disulfide bonds with three experimentally determined structures. Bonds in black text are in agreement, red text indicates disagreement. Dashes indicate disulfides that were absent from the protein construct, NR = not resolved. **C.** The N-terminal portion of AlphaFold HCV-C E1 and the cognate crystal structure (PDB:4UOI).

#### Analysis of the E1E2 interface

Despite the structural divergence between Hepaci, Pegi and Pestivirus E2 proteins, we may expect that they share conserved molecular communications with E1. As described in the main text (Figure 4), the majority of E1E2 interactions occur through a common ‘ancestral’ interface, which, in Hepaci and Pegiviruses, is augmented by an ‘additional’ species-specific interface. To provide detailed insights on these E1E2 contact sites we measured the per residue proximity to partner protein for HCV-C, HCV-P, SPgV and BVDV (i.e. for any given E1 residue what is the shortest distance to E2, and vice versa; Fig. S17). The ancestral interface is composed of four elements: i) packing of the E1 and E2 transmembrane domains alongside the E1 helical hairpin (annotated with a circle on Fig. S17B & C) ii) contact between the conserved beta sheet, which is central to E1, and a transmembrane proximal helix in E2 (triangle), iii) and iv) interactions between the stem of E2 and the bridging loop (hexagon) and transmembrane proximal region (star) of E1. The additional interface involves the extended N-terminal tail of E1, found in Hepaci and Pegiviruses, contacting species-specific loops protruding from E2 (squares). For example, HCV-C variable region 2 (VR2) forms one of these contact sites; the VR2 loop is absent from most other Hepaciviruses.

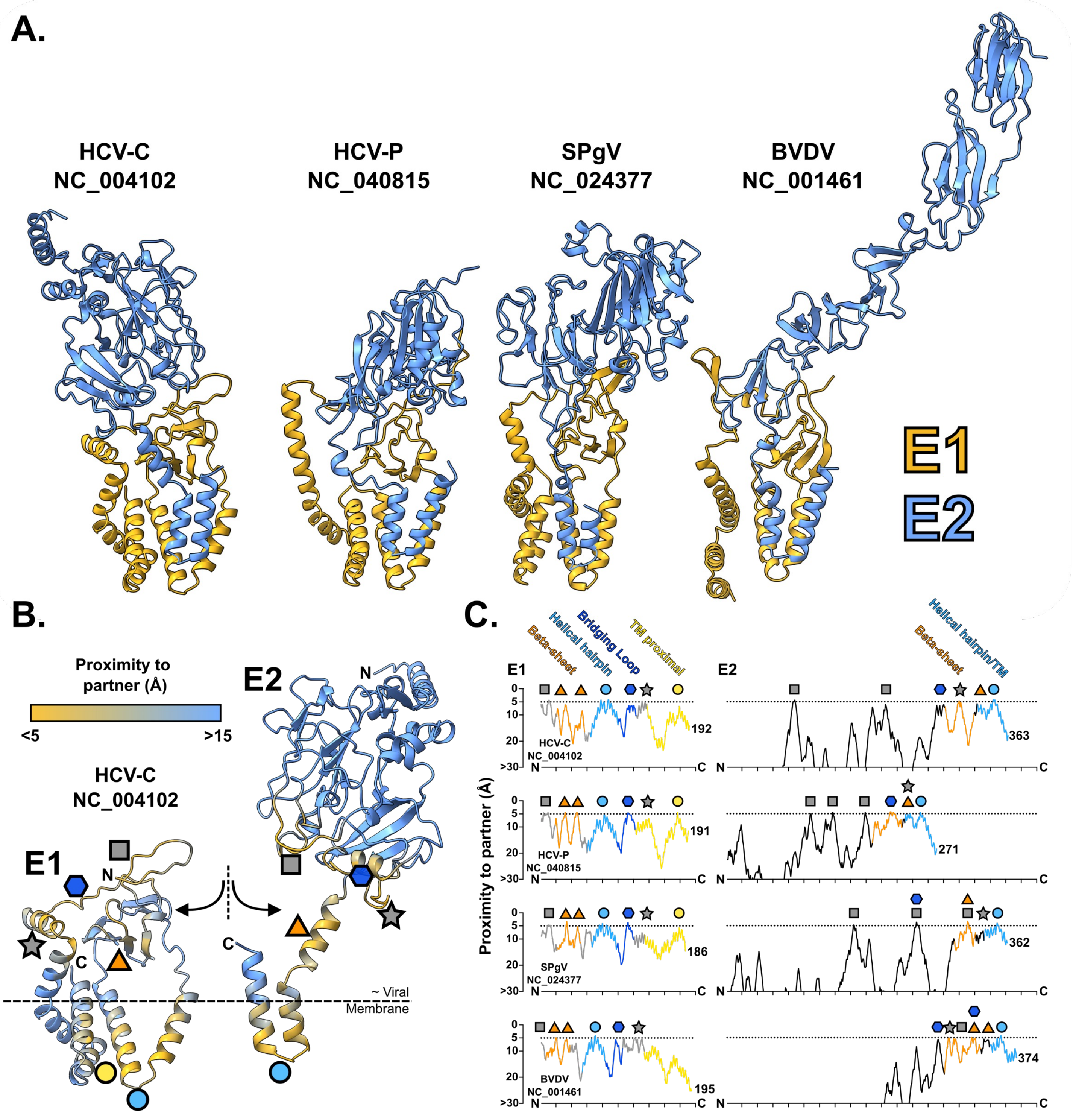


**Figure S17. Comparison of the E1E2 interface in Hepaci, Pegi and Pestiviruses. A.** Ribbon diagrams of the E1E2 complex from HCV-C, HCV-P, SPgV and BVDV. **B.** HCV-C E1E2 complex interaction interface. Residues are colour coded by their shortest distance to the partner protein (Cα to Cα). Symbols annotate contact sites between E1 and E2. The approximate location of the outer leaflet of the viral membrane is inferred by the positions of the E1 and E2 transmembrane domains. **C.** Plots provide E1 and E2 per residue shortest distance to partner protein (i.e. proximity of any given E1 residue to E2, and vice versa) for HCV-C, HCV-P, SPgV and BVDV. Symbols relate to sites of contact, as annotated in B. Lines are colour coded by structural features as in main text Fig. 2A (E1) and Fig. 3C (E2).

### In vitro experimental methods

#### Cells

HEK 293T CD81 KO cells [25] and Huh-7 cells were maintained in Dulbecco's Modified Eagle Medium with L-glutamine supplemented with 10% foetal bovine serum , 1% Penicillin/Streptomycin and 1% non-essential amino acids at 37°C/5% $CO_2$. HEK 293T CD81 KO cells are available upon request.

#### Pseudotyped Virus production

Pseudotyped viruses harbouring HCV-C H77 E1E2 or mutant glycoproteins were produced by co-transfection of approximately 2 million HEK 293T CD81 KO cells using FuGENE6 transfection reagent (Promega) with 1300 ng of an HIV packaging construct (pCMV-dR8.91), 1300 ng of a luciferase reporter plasmid (CSLW), and 100 ng of the given E1E2 expression vector. Pseudotyped virus-containing supernatant was collected at 48 and 72 hours post transfection and pooled. Transfected cells were lysed 72 hours after transfection with 1 mL of cell extraction buffer (Invitrogen) supplemented with protease inhibitor cocktail (Cell Signalling Technology).

#### Mutant E1

Mutant E12 constructs were generated by inserting synthesised sequences (GeneArt, ThermoFisher) containing the mutations of interest into the HCV-H77 E1E2 expression plasmid at XbaI and BstXI restriction sites (parental plasmid is available here: https://www.addgene.org/86983/).

#### Pseudotyped Virus Infection assay

Huh-7 cells were seeded at $3x10^4$ cells per well in a 96 well plate (Greiner) immediately prior to infection with 50µl of pseudotyped virus. Cells were lysed and analysed for luciferase activity using a BrightGlo assay kit (Promega) 48-72 hours post infection.

#### ELISA

Immulon 4 HBX plates (Thermo Scientific) were coated with 0.25 g of Galanthus nivalis lectin (Sigma-Aldrich) in 100µl of Phosphate Buffered Saline (PBS) and incubated overnight at room temperature. Plates were washed 3x with PBS + 0.05% Tween-20 (PBS-T) and blocked with 200µl/well of 2% milk in PBS-T at room temperature for 2 hours. Plates were then washed after blocking 3x with PBS-T. Cell lysates containing E1E2 were added and incubated at room temperature for 2 hours. Following coating with cell lysates, plates were washed 3x and then incubated for 1 hour at room temperature with one of the following: anti-E1 mAb (GeneTex, 0.5µg/mL); anti-E2 mAb (AP33 [26], 0.1µg/mL); anti-E1/E2 conformationally sensitive mAb (AR5A [27], 1µg/mL); or CD81-LEL-hFc (Bio-Techne, 2 µg/mL). After 3 more PBST washes, plates were incubated with either HRP-labelled anti-mouse IgG (1:1000, Sigma-Aldrich) or anti-human IgG (1:10,000, Sigma-Aldrich) for 1 hour at room temperature. Following a final 6 washes with PBST, plates were developed by adding 100µl of TMB substrate (Thermo Scientific). The reaction was stopped by adding 100µl of stop solution (Thermo Scientific). Absorbance was measured at 450 nm. For AR5A ELISAs after plates were incubated with cell lysate, a second blocking step with 200µl of HEK 293T CD81 KO cell lysate was added for 2 hours at room temperature; this step was necessary to eliminate background signal associated with anti-human secondary.
